# A-type lamins anchor emerin at the inner nuclear membrane via two independent binding sites

**DOI:** 10.1101/2025.09.01.673543

**Authors:** Jacob Odell, Kristen Nedza, Jan Lammerding

## Abstract

Lamins form a dense meshwork at the inner surface of the inner nuclear membrane (INM), where they interact with other nuclear envelope proteins such as emerin. Emerin is an integral membrane protein that is part of the LEM (LAP2/emerin/MAN1) domain family, and mutations in either emerin or lamin A/C can result in Emery-Dreifuss muscular dystrophy (EDMD) and other striated muscle diseases. Emerin is retained at the INM through direct interaction with lamin A/C, and emerin’s proper subcellular localization is critical for its ability to influence the mechanical properties of the nucleus and participate in various signaling processes. Nonetheless, the requirements for interaction between emerin and lamin A/C at the INM remain incompletely understood. Here, we report that two distinct regions of lamin A/C are each sufficient to properly localize emerin to the INM and prevent emerin’s lateral diffusion within the INM. In addition to a previously described region of the lamin A/C tail domain able to bind emerin, we identify a novel emerin-interacting domain comprising the linker between the rod and Ig-like fold domains of lamin A/C. We further demonstrate that stably anchoring emerin to the INM requires assembly of A-type lamins into a filamentous network. Collectively, our findings suggest a revised model for emerin retention at the INM, which predicts that two independent lamin A/C domains are required to retain emerin at the nuclear envelope.

## Introduction

Emerin is an INM protein involved in nuclear architecture, gene regulation, and signal transduction (Berk *et al*., 2013). Emerin interacts with structural proteins of the INM, such as lamins and components of the linker of nucleoskeleton and cytoskeleton (LINC) complex (Muchir and Worman, 2007). Mutations in the genes encoding emerin (*EMD*) or lamin A/C (*LMNA*) can result in X-linked and autosomal dominant Emery-Dreifuss muscular dystrophy (EDMD), respectively (Bione *et al*., 1994; Worman, 2012; Koch and Holaska, 2014). Emerin, which is part of a family of proteins containing the LEM (LAP2 (lamina-associated polypeptide 2)/emerin/MAN1) domain, is a type-II integral membrane protein with an N-terminal nucleoplasmic domain, a small (11-residue) luminal domain at its C-terminus, and its transmembrane domain near the C-terminus (Berk *et al*., 2013). Given its small size, emerin can easily diffuse between the INM and the outer nuclear membrane and endoplasmic reticulum (ER), which form a continuous membrane system. However, in many cell types, emerin is anchored at the INM through interactions with lamin A/C, corresponding to a diffusion-retention model (Sullivan *et al*., 1999; Östlund *et al*., 2006; Zwerger *et al*., 2015). Deletion of *Lmna* in mice results in mislocalization of many nuclear envelope proteins including emerin, which is displaced from the INM to the ER in muscle cells and fibroblasts, leading to a muscular dystrophy phenotype that mimics the disease presentation of EDMD in humans (Sullivan *et al*., 1999; Holt *et al*., 2003). Similarly, *LMNA* mutations frequently lead to mislocalization of emerin from the INM in cells from patients with EDMD (Raharjo *et al*., 2001), suggesting that impaired interaction between Lamin A/C and emerin contributes to the disease pathogenesis. Of note, however, not all EDMD-causing *LMNA* mutations lead to mislocalization of emerin from the nuclear envelope (Reichart *et al*., 2004), and although the *LMNA* regions coding for coil 2 and the Ig-like fold of Lamin A (LaA) and C (LaC) are hotspots of mutations (Bertrand *et al*., 2020), EDMD-causing mutations are found along the length of the *LMNA* gene (Worman, 2012). Thus, the contribution of different LaA/C protein domains to emerin interaction in intact cells remains incompletely understood. In addition to the A-type lamins (primarily LaA and LaC), mammalian cells also express two B-type lamins, lamin B1 (*LMNB1*) and lamin B2 (*LMNB2*), but the interaction between lamins and emerin is specific to A-type lamins and does not extend to B-type lamins. Consequently, emerin is mislocalized in many tissues in *Lmna*^−/−^ mice, despite the maintained presence of both B-type lamins (Sullivan *et al*., 1999). Yet, it is unknown what the key differences are between the A-type and B-type lamins that enable LaA/C to interact with emerin and preclude interaction between lamin B1 (LaB1) and emerin.

Several efforts have been made to elucidate details of the interaction between A-type lamins and emerin using in vitro approaches. Lamins possess a tripartite domain structure with a central coiled-coil rod domain flanked by a small N-terminal head domain and a C-terminal tail that contains a nuclear localization signal (NLS) and an immunoglobulin-like (Ig-like) fold (Aebi *et al*., 1986; Buchwalter, 2023). A previous yeast-two-hybrid screen indicated that the C-terminal tail of LaA (aa 385-566) was sufficient to bind emerin (Sakaki *et al*., 2001). Further work using recombinant LaA and emerin identified a direct interaction between full-length LaA and emerin using surface plasmon resonance (Clements *et al*., 2000) and confirmed binding of the LaA tail (aa 385-646) and emerin (Berk *et al*., 2014), although atomic-level details of direct interaction between LaA and emerin have not yet been revealed. A recent crystal structure describes a ternary complex involving the LaA Ig-like fold (aa 411-566) binding to barrier-to-autointegration factor (BAF), which in turn binds to emerin, thus providing a mechanism for an indirect interaction between LaA and emerin involving the LaA tail domain (Samson *et al*., 2018). Collectively, these results indicate that the LaA tail domain is required for the interaction between LaA and emerin. However, previous studies investigating the interaction between lamins and emerin in cells have been restricted to expressing wild-type or mutant LaA/C, without examining the role of specific domains or differences between A-type and B-type lamins (Raharjo *et al*., 2001; Zwerger *et al*., 2013; Odell *et al*., 2024).

To better define which regions of A-type lamins specifically contribute to the interaction with emerin in living cells, we generated a series of expression constructs to allow for expression of full-length lamins, truncated lamins, and chimeric lamins under the control of a doxycycline-inducible promoter. Following expression of these constructs in *Lmna*^−/−^ MEFs or cells lacking all lamins, we determined the ability of each construct to restore the proper nuclear localization of endogenous emerin and restrict the mobility of GFP-emerin at the INM. This rescue approach in living cells overcomes the limitation of previous in vitro binding assays, which cannot fully capture the physiological interaction of LaA/C and emerin in live cells, including the assembly of lamins into a network and the role of the nuclear membranes, which are likely important for the physiological interaction between A-type lamins and emerin. Using our novel experimental system, we identified a previously overlooked region of LaA/C comprising the linker between the rod and Ig-like fold domains that is sufficient for LaA/C to interact with emerin and anchor it to the INM. When this region was inserted into LaB1, which does not normally interact with emerin, it conferred emerin binding ability to LaB1. Additionally, we found that the formation of a lamin network at the nuclear periphery is required to anchor emerin at the INM, as lamin constructs lacking the rod domain, which is essential for filament assembly, or expression of engineered proteins that prevent the incorporation of LaA/C into the nuclear lamina, failed to anchor emerin to the INM. Collectively, our work defines the molecular requirements to tightly anchor emerin at the INM, including the additive contributions of two independent LaA/C emerin-interacting domains and the formation of a stable A-type lamin network.

## Results

### Localization and anchorage of emerin to the INM requires A-type lamins

Emerin is anchored in place at the INM in many cells, including fibroblasts, through interactions with LaA/C (Figure 1) (Sullivan *et al*., 1999; Vaughan *et al*., 2001; Östlund *et al*., 2006). In contrast to wild-type cells, *Lmna*^−/−^ MEFs display abnormal emerin subcellular localization, with emerin no longer concentrated in the nucleus and instead enriched in the ER (Figure 1A-B) (Sullivan *et al*., 1999; Vaughan *et al*., 2001; Östlund *et al*., 2006). To define the contribution of each lamin isoform and lamin domain to this emerin interaction, we designed a panel of exogenous expression constructs to introduce specific full-length or modified human lamin isoforms into *Lmna*^−/−^ MEFs at physiological expression levels (Figure 1C) (Odell *et al*., 2025). Expression of these constructs is driven by a doxycycline (dox) inducible promoter, which allows for a comparison of emerin localization at baseline (i.e., in the absence of exogenous lamin, “no dox”) versus after doxycycline-induced lamin expression. As expected, emerin was mislocalized in all “no dox” conditions relative to wild-type MEFs, matching the phenotype of *Lmna*^−/−^ MEFs (Supplemental Figure 1).

**Figure 1:**
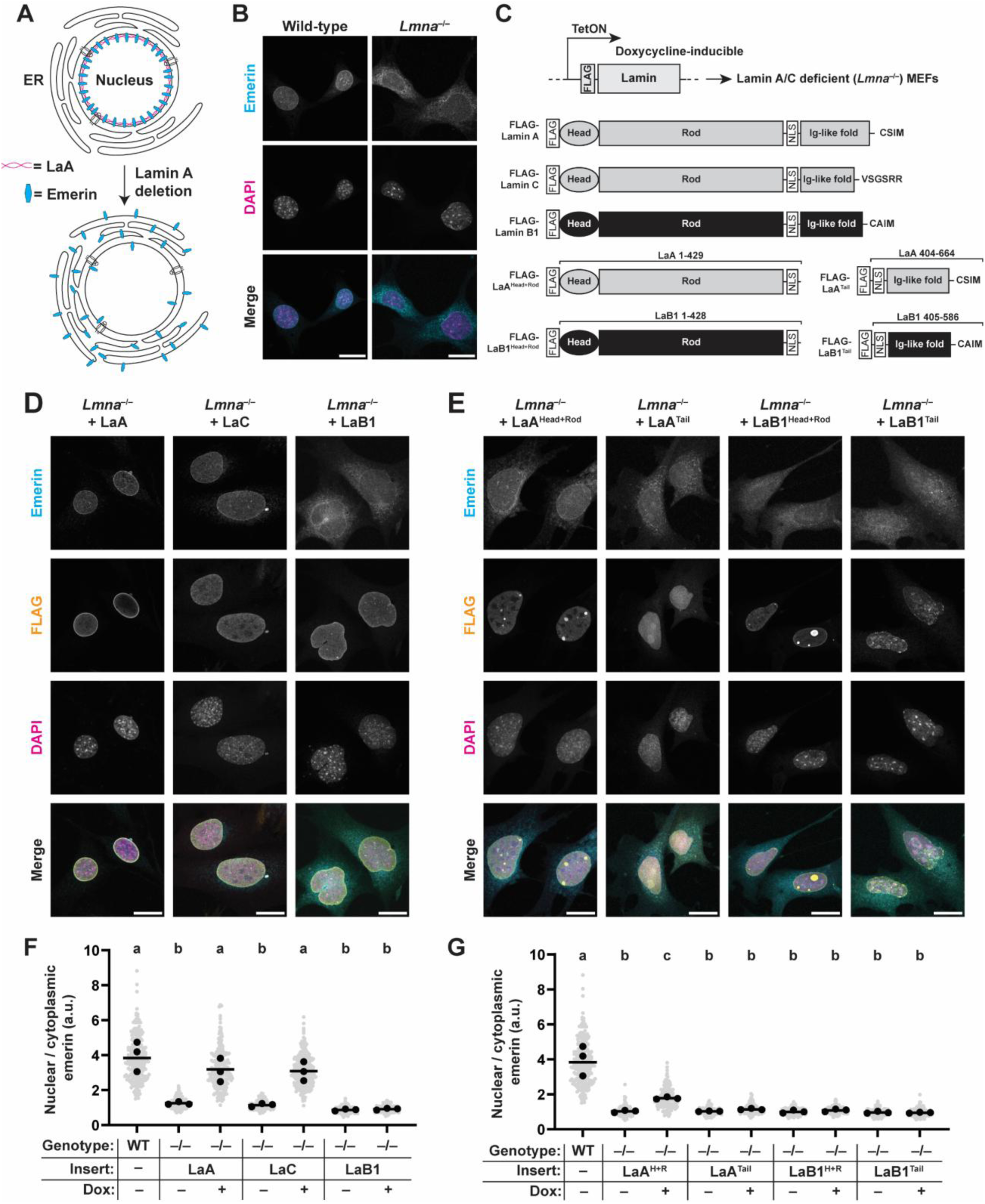
Nuclear localization of emerin in mouse embryo fibroblasts is dependent on A-type lamins. (**A**) Schematic depiction of the diffusion-retention model of emerin anchoring to the INM by A-type lamins. Following loss of A-type lamins, emerin is no longer anchored at the INM and diffuses to the outer nuclear membrane and the endoplasmic reticulum, which form a continuous membrane system with the INM. (**B**) Representative micrographs of wild-type and *Lmna*^−/−^ MEFs immunofluorescently labeled for endogenous emerin. Scale bar = 20 µm. (**C**) Exogenous expression strategy to introduce a lamin construct of interest under the control of a doxycycline-inducible promoter. All constructs possess a small N-terminal FLAG tag, allowing for detection by anti-FLAG antibodies. For truncated constructs, the residues derived from the respective full-length proteins are indicated. (**D**) Immunofluorescence labeling for emerin in *Lmna*^−/−^ MEFs expressing full-length LaA, LaC, or LaB1. FLAG channel depicts the localization of the exogenous lamin. Scale bar = 20 µm. (**E**) Immunofluorescence labeling for emerin in *Lmna*^−/−^ MEFs expressing lamin truncations. Scale bar = 20 µm. (**F-G**) Quantification of mean nuclear/cytoplasmic fluorescence intensity of immunofluorescently labeled endogenous emerin in *Lmna*^−/−^ MEFs expressing full-length lamins (**F**) or truncated lamins (**G**). Grey points indicate measurements from individual cells, black points indicate replicate means, and bars indicate overall means. Sets of points with the same letter above them are not significantly different, whereas different letters indicate *p* < 0.05, based on one-way ANOVA with Tukey’s multiple comparison test.

Dox-induced expression of LaA or LaC significantly improved nuclear localization of emerin in *Lmna*^−/−^ MEFs (Figure 1D), consistent with prior reports that reintroduction of A-type lamins can rescue emerin localization in these cells (Sullivan *et al*., 1999; Östlund *et al*., 2006; Odell *et al*., 2024). In contrast, overexpression of LaB1 did not affect emerin localization (Figure 1D). We did not observe any difference in the ability of LaA vs LaC to rescue the subcellular localization of emerin, suggesting that both A-type lamins possess similar abilities to interact with emerin (Figure 1F).

We next assessed the subcellular localization of emerin in *Lmna*^−/−^ MEFs expressing truncated lamin constructs consisting of either the head, rod, and NLS of LaA, but lacking the Ig-like fold (LaA^Head+Rod^), or containing the NLS and Ig-like fold of LaA but lacking the head and rod domains (LaA^Tail^). Despite lacking the LaA tail, expression of the LaA^Head+Rod^ construct improved the nuclear localization of endogenous emerin in *Lmna*^−/−^ MEFs, albeit to a lesser extent than full-length LaA (Figure 1E). This surprising result suggests that the interaction between LaA and emerin is mediated, at least in part, by residues found in LaA^Head+Rod^. In contrast to the LaA^Head+Rod^ construct, expression of LaA^Tail^ had no effect on emerin localization in lamin-deficient cells (Figure 1E). Of note, due to the absence of the rod domain, LaA^Tail^ cannot form dimers or higher-order filaments at the nuclear lamina but is instead present as a soluble protein within the nucleoplasm (Heitlinger *et al*., 1992) (Odell *et al*., 2025). Neither of the corresponding truncations of LaB1 significantly altered the subcellular localization of emerin (Figure 1E, G), consistent with our findings with full-length LaB1 and confirming that the proper localization of emerin is dependent on A-type lamins.

To eliminate the possibility that the LaB1 expression constructs may be interacting or competing with endogenous B-type lamins in *Lmna*^−/−^ MEFs, which might mask their ability to interact with emerin, we expressed full-length lamins and truncations in triple-lamin-knockout (TKO: *Lmna*^−/−^, *Lmnb1*^−/−^, *Lmnb2*^−/−^) MEFs and assessed emerin subcellular localization. The results in TKO MEFs closely mirrored those obtained using *Lmna*^−/−^ MEFs: expression of full length LaA and LaC, but not LaB1, rescued emerin nuclear localization, and the only lamin truncation construct that affected emerin localization was LaA^H+R^ (Supplemental Figure 2A-B). These results highlight that the interaction between lamins and emerin is specific to the A-type and not the B-type lamins.

As an additional readout for interaction between lamins and emerin, we performed fluorescence recovery after photobleaching (FRAP) experiments using GFP-emerin transiently expressed in *Lmna*^−/−^ MEFs. FRAP involves photobleaching fluorescently tagged proteins in a defined region of the cell and monitoring the recovery of fluorescence intensity within the bleached region, which provides information about the lateral mobility of the tagged protein (Reits and Neefjes, 2001). Consistent with the idea that A-type lamins anchor emerin in place at the INM, previous work has demonstrated an increased recovery rate of GFP-emerin following photobleaching in *Lmna*^−/−^ cells compared to wild-type MEFs (Östlund *et al*., 2006). Confirming the findings from this prior study, we observed a significant increase in the recovery of GFP-emerin following photobleaching of the nuclear envelope in *Lmna*^−/−^ MEFs compared to wild-type controls (Figure 2A). Expression of either LaA or LaC reduced the mobility of GFP-emerin in *Lmna*^−/−^ MEFs to wild-type levels (Figure 2B-C). We did not find any difference in the rescue achieved by LaA versus LaC, in agreement with the immunofluorescence localization data for endogenous emerin (Figure 1F) and consistent with the idea that both A-type lamins can equally interact with emerin and restrict its mobility at the INM. Overexpression of LaB1 in *Lmna*^−/−^ MEFs failed to rescue the mobility of GFP-emerin (Figure 2D), further emphasizing that lamin interaction with emerin is specific to A-type lamins. Quantification of the median recovery half-time (*t*_½_) of GFP-emerin after photobleaching revealed a significant reduction in the recovery time of GFP-emerin following loss of A-type lamins, which was completely restored to wild-type levels following reintroduction of LaA or LaC but remained unchanged by overexpression of LaB1 in the *Lmna*^−/−^ MEFs (Figure 2E).

**Figure 2:**
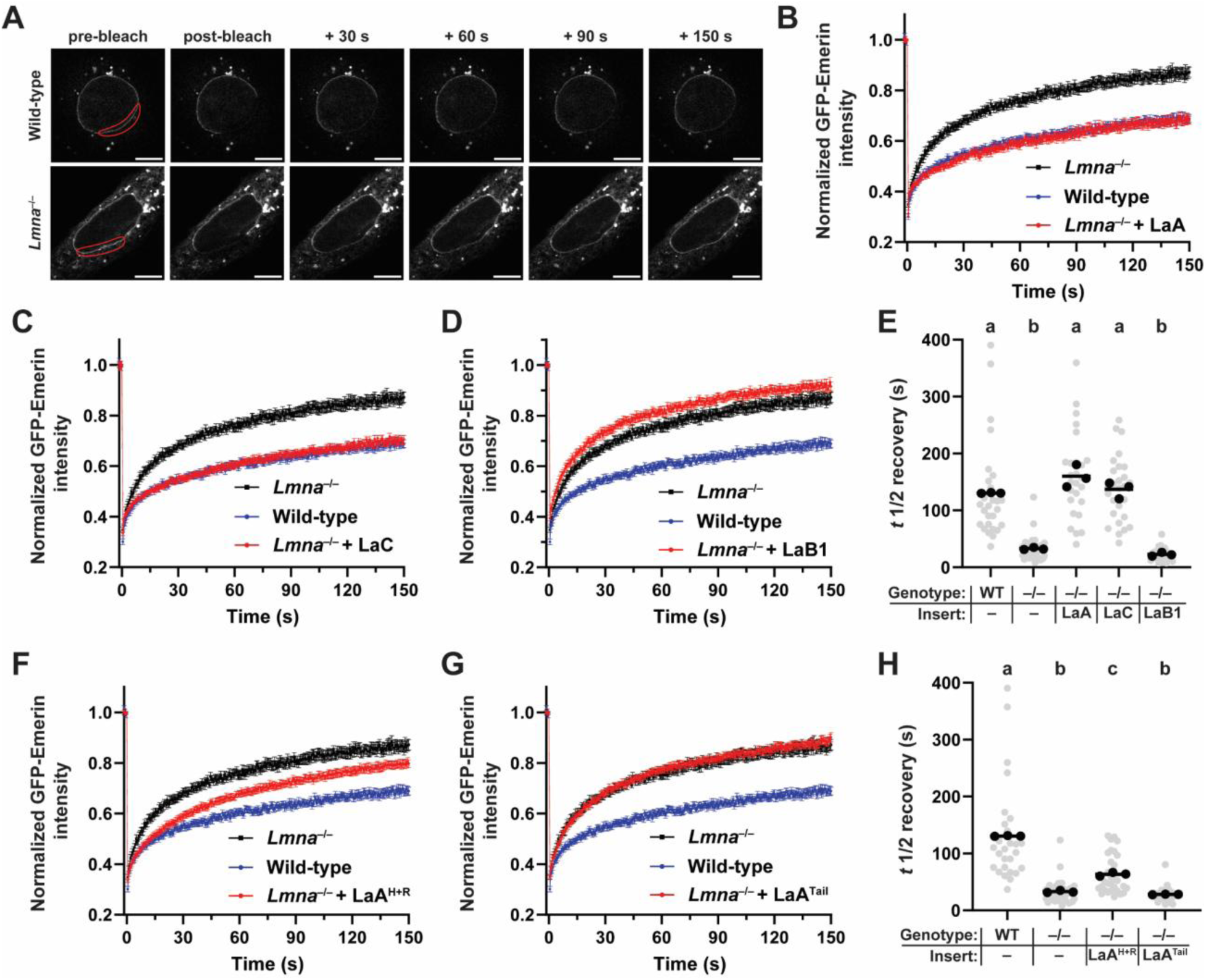
Mobility of GFP-emerin at the nuclear envelope is dependent on A-type lamins and partially rescued by LaA^Head+Rod^. (**A**) Representative frames of a time series showing FRAP experiment on GFP-emerin transiently expressed in wild-type or *Lmna*^−/−^ MEFs. The target bleach area is outlined in red, and the time in seconds following bleach is indicated above each set of images. Scale bar = 10 µm. (**B**) Recovery curves of GFP-emerin following photobleaching in wild-type, *Lmna*^−/−^, and LaA rescue cells. Mean fluorescence intensity within the bleached region was normalized to the prebleach intensity, and corrections were made to account for photobleaching resulting from continuous imaging. Each point represents the average of 25-30 cells, bars indicate s.e.m. (**C-D**) Recovery curves of GFP-emerin following photobleaching in *Lmna*^−/−^ MEFs expressing LaC (**C**) or LaB1 (**D**). Data from wild-type and *Lmna*^−/−^ cells are added for reference. (**E**) Quantification of *t*_½_ median recovery for the data presented in panels B-D. Lower values for *t*_½_ recovery indicate faster rates of recovery and more mobile GFP-emerin. (**F-H**) Data from FRAP experiments performed on *Lmna*^−/−^ MEFs expressing LaA^Head+Rod^ (**F**) or LaA^Tail^ (**G**). Grey points indicate measurements from individual cells, black points indicate replicate means, and bars indicate overall means. Sets of points with the same letter above them are not significantly different, whereas different letters indicate *p* < 0.05, based on one-way ANOVA with Tukey’s multiple comparison test.

FRAP experiments on *Lmna*^−/−^ MEFs expressing the LaA truncations further corroborated our conclusions from the immunofluorescence labeling results (Figure 1E, G). Expression of LaA^Head+Rod^ partially reduced the mobility of GFP-emerin at the INM (Figure 2F), whereas LaA^Tail^ did not affect the mobility of GFP-emerin (Figure 2G) in *Lmna*^−/−^ MEFs. Accordingly, expression of LaA^Head+Rod^, but not LaA^Tail^, significantly increased the median *t*_½_ recovery relative to baseline *Lmna*^−/−^ cells (Figure 2H). Collectively, these results support our proposition that LaA residues contained within LaA^Head+Rod^ (aa 1-429) interact with emerin, as this construct was sufficient to (partially) restore the proper subcellular distribution and mobility of emerin in *Lmna*^−/−^ cells.

To determine whether other LEM domain proteins that interact with LaA are anchored by LaA similar to emerin, we performed FRAP experiments on wild-type and *Lmna*^−/−^ MEFs expressing Man1-GFP (Supplemental Figure 3A-B). We did not observe any significant difference in Man1-GFP mobility between wild-type and *Lmna*^−/−^ MEFs (Supplemental Figure 3B-D). Moreover, re-introduction of LaA in *Lmna*^−/−^ MEFs did not significantly alter the mobility of Man1-GFP (Supplemental Figure 3C-D), indicating that unlike emerin, Man1 does not require interaction with A-type lamins to anchor it at the nuclear membranes. These results are largely consistent with a previous study (Östlund *et al*., 2006), which, despite finding subtle differences in Man1 localization between *Lmna*^−/−^ and wild-type MEFs, observed only minimal differences in Man1-GFP mobility between these cells, in contrast to emerin, which showed a substantial increase in mobility at the nuclear envelope in the *Lmna*^−/−^ MEFs (Östlund et al., 2006). These results suggest that the interaction between emerin and LaA is not mediated exclusively by the LEM domain or BAF binding, both of which are shared by Man1.

### The LaA head domain is not sufficient to drive nuclear localization of emerin

We next aimed to determine which regions within LaA^Head+Rod^ were crucial for its ability to interact with emerin, as prior work had focused mainly on interactions involving residues in the LaA Ig-like fold (Sakaki *et al*., 2001; Berk *et al*., 2014; Samson *et al*., 2018). To identify candidate regions within LaA^Head+Rod^ that bind to emerin, we performed a sequence alignment of human LaA and LaB1 (Supplemental Figure 4) and searched for regions that are present in the LaA^Head+Rod^ construct, which interacts with emerin (Figure 1G), and that are divergent from LaB1^Head+Rod^, which does not interact with emerin (Figure 1G). Since the head domains of LaA and LaB1 (aa 1-33 or 1-34) fit both these criteria, we considered whether the LaA-specific head domain may drive interaction with emerin. To test this idea, we generated a series of head-swapped lamin constructs that combined the head of one lamin paralog with the remainder of the other, as well as head-swapped truncations that fused the head of one paralog with the rod and NLS of the other (Figure 3A). If the LaA head domain were responsible for the ability of LaA to interact with emerin, then we would expect that the LaA^Head^-B1^Rod+Tail^ and LaA^Head^-B1^Rod^ constructs would restore the proper nuclear localization of emerin. However, this was not the case (Figure 3C-E). The addition of the LaA head did not confer emerin-binding ability to LaA^Head^-B1^Rod+Tail^ or LaA^Head^-B1^Rod^ (Figure 3C, E). Similarly, the substitution of the LaB1 head in place of the LaA head did not impair the ability of LaB1^Head^-A^Rod+Tail^ or LaB1^Head^-A^Rod^ to properly rescue the subcellular nuclear localization of emerin (Figure 3B, D). These data demonstrate that the ability of LaA to interact with emerin is not determined by the LaA head domain.

**Figure 3:**
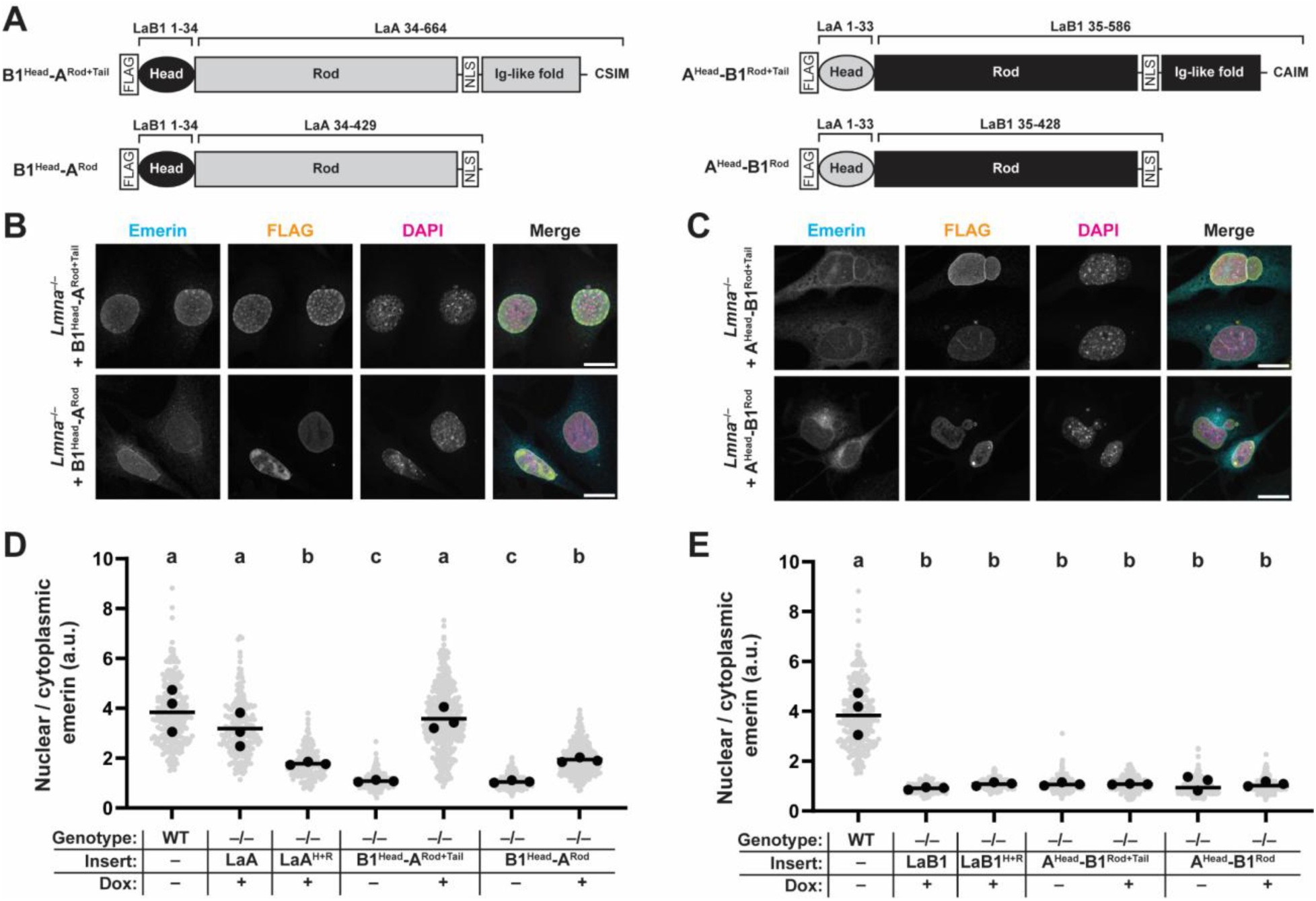
The ability of lamins to interact with emerin is not determined by the lamin head domain. (**A**) Schematic representation of “head-swapped” lamin expression constructs. Residues from LaA versus LaB1 are indicated above each construct. Similar to other constructs used in this study, all head-swapped lamin constructs contain an N-terminal FLAG tag for detection of exogenous lamin by immunofluorescence. (**B-C**) Representative immunofluorescence images of *Lmna*^−/−^ MEFs expressing the head-swapped lamin constructs. FLAG signal indicates the expression of the respective head-swapped lamin protein. Scale bar = 20 µm. (**D-E**) Quantification of the mean nuclear-to-cytoplasmic fluorescence intensity ratio of endogenous emerin in *Lmna*^−/−^ MEFs expressing head-swapped lamin constructs. The rescue achieved by the ‘wild-type’ lamins is shown for each plot for comparison purposes to demonstrate that the substitution of the head domain does not alter the ability of each construct to influence the subcellular localization of emerin. Grey points indicate measurements from individual cells, black points indicate replicate means, and horizontal bars indicate overall means. Sets of points with the same letter above them are not significantly different, whereas different letters indicate *p* < 0.05, based on one-way ANOVA with Tukey’s multiple comparison test.

### The LaA linker between rod and Ig-like fold domains comprises a novel emerin-interacting domain

We identified an additional divergent region of LaA and LaB1 that met our criteria, consisting of 25 amino acids located in the flexible linker (L) between the end of coil 2b of the rod domain and the Ig-like fold (Figure 4A). The lamin linker region, which contains the NLS, is highly conserved among LaB1 homologs (Dou *et al*., 2015) and differs between LaA and LaB1 in several vertebrates, including mouse and humans (Supplemental Figure 4). We hypothesized that this region might comprise a not previously recognized emerin-interacting domain of LaA (Figure 4A).

**Figure 4:**
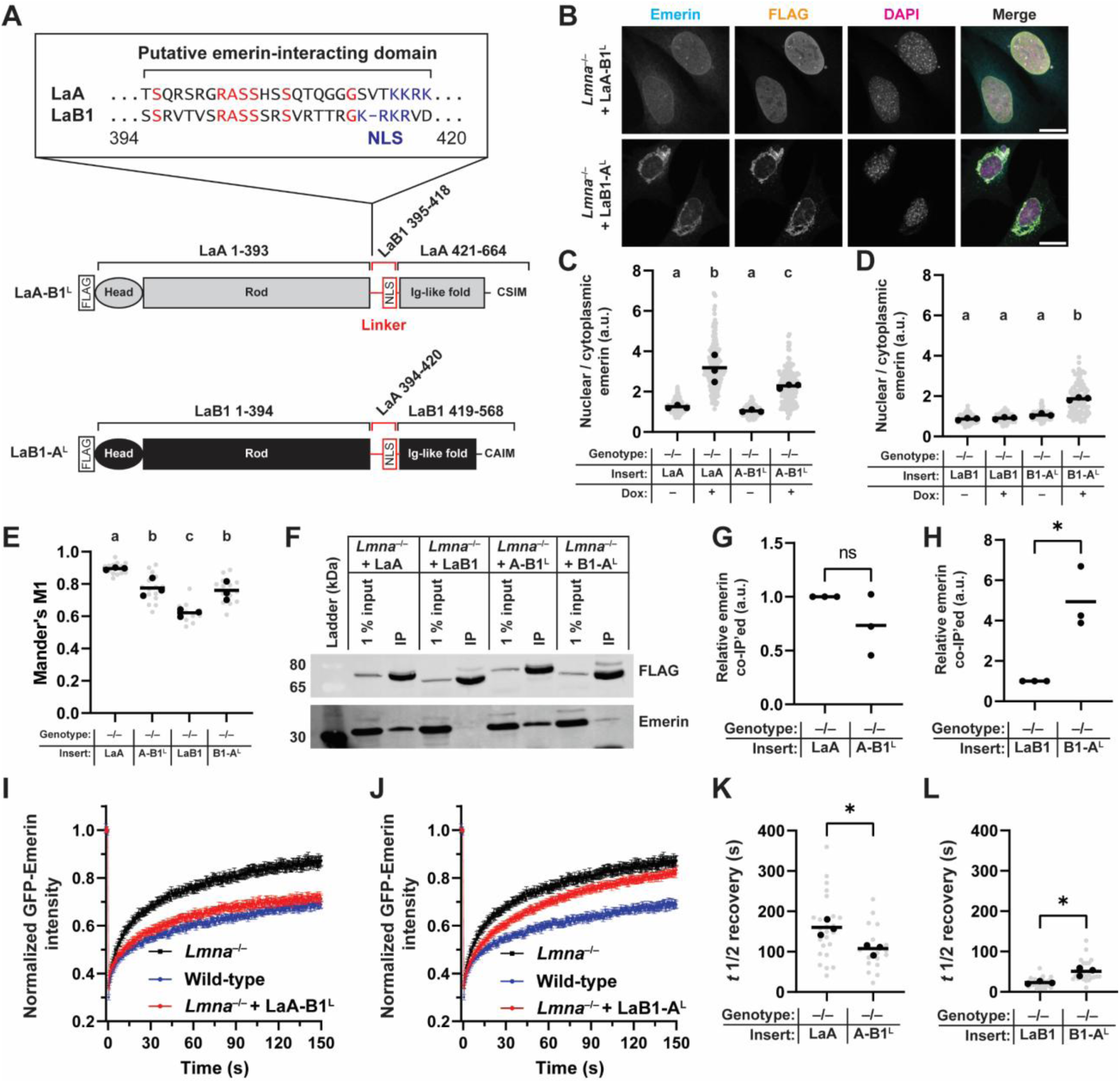
Identification of a novel emerin-interacting domain of LaA. (**A**) Schematic of the putative emerin-interacting domain of LaA and alignment between the corresponding regions of LaA and LaB1. Residues colored in red are identical between the lamin isoforms; the NLS is denoted in blue. (**B**) Representative immunofluorescence images of *Lmna*^−/−^ MEFs expressing the Linker-swapped lamins. FLAG signal indicates the localization of the exogenous lamin construct. Scale bar = 20 µm. (**C-D**) Quantification of mean nuclear-to-cytoplasmic fluorescence intensity ratio of endogenous emerin in *Lmna*^−/−^ MEFs expressing the indicated lamin constructs. The rescue achieved by the ‘wild-type’ LaA (**C**) or LaB1 (**D**) is shown for each plot for comparison. Grey points indicate measurements from individual cells, black points indicate replicate means, and bars indicate overall means. (**E**) Colocalization analysis comparing the colocalization between emerin and FLAG signal for wild-type LaA and LaB1 versus Linker-swapped constructs. Mander’s M1 represents the proportion of pixels in the emerin channel that overlap with pixels in the FLAG channel, with a maximum value of 1 for perfect correlation. Grey points indicate measurements from separate fields of view, black points indicate replicate means, and bars indicate overall means. For C-E, sets of points with the same letter above them are not significantly different, whereas different letters indicate *p* < 0.05, One-way ANOVA with Tukey’s multiple comparison test. (**F**) Immunoblot of co-immunoprecipitation experiments performed using wild-type or Linker-swapped lamins as bait. IP was performed using an anti-FLAG antibody that recognized the FLAG epitope present on all expression constructs. 1% input was set aside prior to IP to show equal amounts of starting material. (**G-H**) Quantification of emerin that co-IP’ed with the indicated constructs. Points indicate values from 3 independent co-IP experiments. (**I-J**) FRAP curves of mean GFP-emerin intensity in *Lmna*^−/−^ MEFs expressing LaA-B1^L^ (**I**) or LaB1-A^L^ (**J**). (**K-L**) Quantification of *t*_½_ recovery of GFP-emerin following photobleaching in *Lmna*^−/−^ MEFs expressing Linker-swapped lamins. Grey points indicate measurements from individual cells, black points indicate replicate means, and bars indicate overall means. For G, H, K, and L, sets of points with the same letter above them are not significantly different, whereas different letters indicate *p* < 0.05, based on unpaired two-tailed *t*-tests.

In this case, substituting these LaA residues in place of the corresponding region of LaB1 (LaB1 - A^L^) should be sufficient to confer emerin-binding abilities to LaB1. Conversely, the replacement of these residues from LaA with the corresponding divergent sequence from LaB1 (LaA -B1^L^) would be expected to reduce, but not completely eliminate, the ability of LaA to interact with emerin, given the presence of a second emerin-interacting domain within the Ig-like fold of A-type lamins (Sakaki *et al*., 2001; Berk *et al*., 2014; Samson *et al*., 2018). To test this hypothesis, we generated expression constructs where the linker residues were swapped between LaA and LaB1, referred to as LaB1-A^L^ and LaA -B1^L^, respectively (Figure 4A). Both constructs localized to the nucleus (Figure 4B-C). Expression of LaB1 -A^L^ in *Lmna*^−/−^ MEFs improved nuclear localization of endogenous emerin compared to *Lmna*^−/−^ MEFs (Figure 4B-C), with a similar magnitude of rescue as that observed with LaA^Head+Rod^ (Figure 1F), consistent with our prediction. Expression of the LaA-B1^L^ construct led to a slight reduction in localization of emerin to the nucleus compared with expression of wild-type LaA but still improved nuclear localization of emerin compared to *Lmna*^−/−^ MEFs not expressing exogenous LaA (Figure 4B-C). Taken together, these data suggest that the 25 amino acids from LaA within the linker region between the rod and Ig-like fold contains an independent A-type lamin emerin-interaction domain.

The LaB1-A^L^ construct had an unexpected spatial distribution, as it was partially enriched in a thick layer around the nucleus, possibly in the ER membrane (Figure 4B). Since emerin was co-enriched in these same areas (bottom panel of Figure 4B), we performed quantitative colocalization analysis between the FLAG signal and the emerin signal on *Lmna*^−/−^ MEFs expressing wild-type lamins or the linker-swapped constructs. Full-length LaA exhibited the greatest colocalization with emerin, indicated by a larger Mander’s M1 colocalization index. In contrast, LaB1 had the lowest colocalization with emerin (Figure 4E). Removal of the LaA-specific linker via the LaA -B1^L^ construct reduced the colocalization with emerin relative to wild-type LaA, consistent with the incomplete rescue from this construct on the nuclear localization of emerin. Conversely, the LaB1 -A^L^ construct had improved colocalization with emerin relative to wild-type LaB1 (Figure 4E), further suggesting that the LaA linker residues are involved in interaction with emerin.

As an independent method to probe for the interaction between lamins and emerin, we performed co-immunoprecipitation (co-IP) experiments using full-length LaA and LaB1 or the linker-swapped constructs. Each of the exogenous lamin constructs was pulled down via its N-terminal FLAG tag (Figure 4F). Wild-type, full-length LaA had the greatest ability to co-IP emerin, whereas the co-IP was weakest for LaB1 (Figure 4F-H). Whereas the LaA -B1^L^ chimeric construct was not significantly impaired in its ability to co-IP emerin (Figure 4G), likely due to its second emerin-interacting domain within the LaA Ig-like fold, the LaB1-A^L^ construct had a significantly improved ability to co-IP emerin compared to wild-type LaB1 (Figure 4H), suggesting that the LaA linker region is sufficient to confer some emerin-binding ability to LaB1.

To determine whether the LaA linker was sufficient to anchor emerin to the INM, we performed FRAP experiments on *Lmna*^−/−^ cells expressing the linker-swapped constructs. Expression of LaA-B1^L^ in *Lmna*^−/−^ MEFs decreased the mobility of GFP-emerin following photobleaching (Figure 4I), although the rescue achieved by this construct was slightly less than that of wild-type LaA (Figure 4K). Intriguingly, expression of the LaB1-A^L^ construct was sufficient to partially restrict the mobility of GFP-emerin after photobleaching compared to *Lmna*^−/−^ MEFs (Figure 4J) and increased the *t*_½_ recovery time compared to expression of LaB1 (Figure 4L). Collectively, these results suggest that amino acids 394-420 from LaA are sufficient to confer emerin-binding activity to LaB1 and constitute a previously overlooked LaA emerin-interacting domain.

### The LaA tail requires the formation of a LaA network at the nuclear periphery to retain emerin at the INM

We decided to revisit the LaA^Tail^ construct, which included both regions known to bind emerin but still failed to restore emerin localization in vivo (Figure 1E, G). We hypothesized that LaA^Tail^ failed to rescue emerin localization in cells because it was unable to form a stable structure at the INM. An assembled lamina has been proposed as a prerequisite for interaction of LaA with emerin (Holt *et al*., 2003), leading to the following predictions: (1) Chimeric lamins that possess a rod domain to allow dimerization and one or the other LaA domain able to interact with emerin (LaA linker or LaA^Tail^) should be able to affect emerin localization and anchoring in cells; (2) strengthening the LaA network, for example, by tightly anchoring LaA to the INM, should enhance the interaction with emerin; and (3) disruptions to the assembly of the LaA/C network should abolish the ability to anchor emerin to the INM, even if A-type lamins are present in cells.

To test these predictions, we first generated chimeric lamin constructs that combined the head and rod of LaB1 with the tail of LaA (LaB1^Head+Rod^-A^Tail^), and a reciprocal chimera that fused the head and rod of LaA with the tail of LaB1 (LaA^Head+Rod^-B1^Tail^) (Figure 5A). Note that the LaA^Head+Rod^-B1^Tail^ chimera possesses the LaA-specific linker region capable of interacting with emerin (Figure 4). When expressed in *Lmna*^−/−^ MEFs, both chimeric lamins partially improved the nuclear localization of endogenous emerin (Figure 5B-C) and the same effect was seen when chimeric lamins were expressed in TKO MEFs (Supplemental Figure 2C). In contrast to full-length A-type lamin constructs, though, neither chimeric lamin achieved wild-type levels of nuclear emerin localization (compare Figure 5C with Figure 1F), indicating that they have additive effects. Both chimeras successfully reduced the mobility of GFP-emerin following photobleaching (Figure 5D-E) and partially increased the recovery time of GFP-emerin relative to *Lmna*^−/−^ MEFs (Figure 5F), indicating successful diffusion-retention of emerin at the nuclear membrane. However, neither chimera rescued the mobility or recovery time of GFP-emerin to the same extent as full-length LaA or LaC expressed in *Lmna*^−/−^ MEFs (compare Figure 5D-F with Figure 2B-C). These data suggest that while the LaA tail domain can interact with emerin in vitro, for this domain to affect the subcellular localization or mobility of emerin in vivo, it must be part of a filamentous structure at the nuclear periphery, which requires the lamin head and rod domain. The incomplete rescue of emerin localization and mobility with the chimeric lamin constructs further suggests that the multiple domains of LaA that interact with emerin act additively and that both are required to achieve the full effect on emerin seen for wild-type A-type lamins.

**Figure 5:**
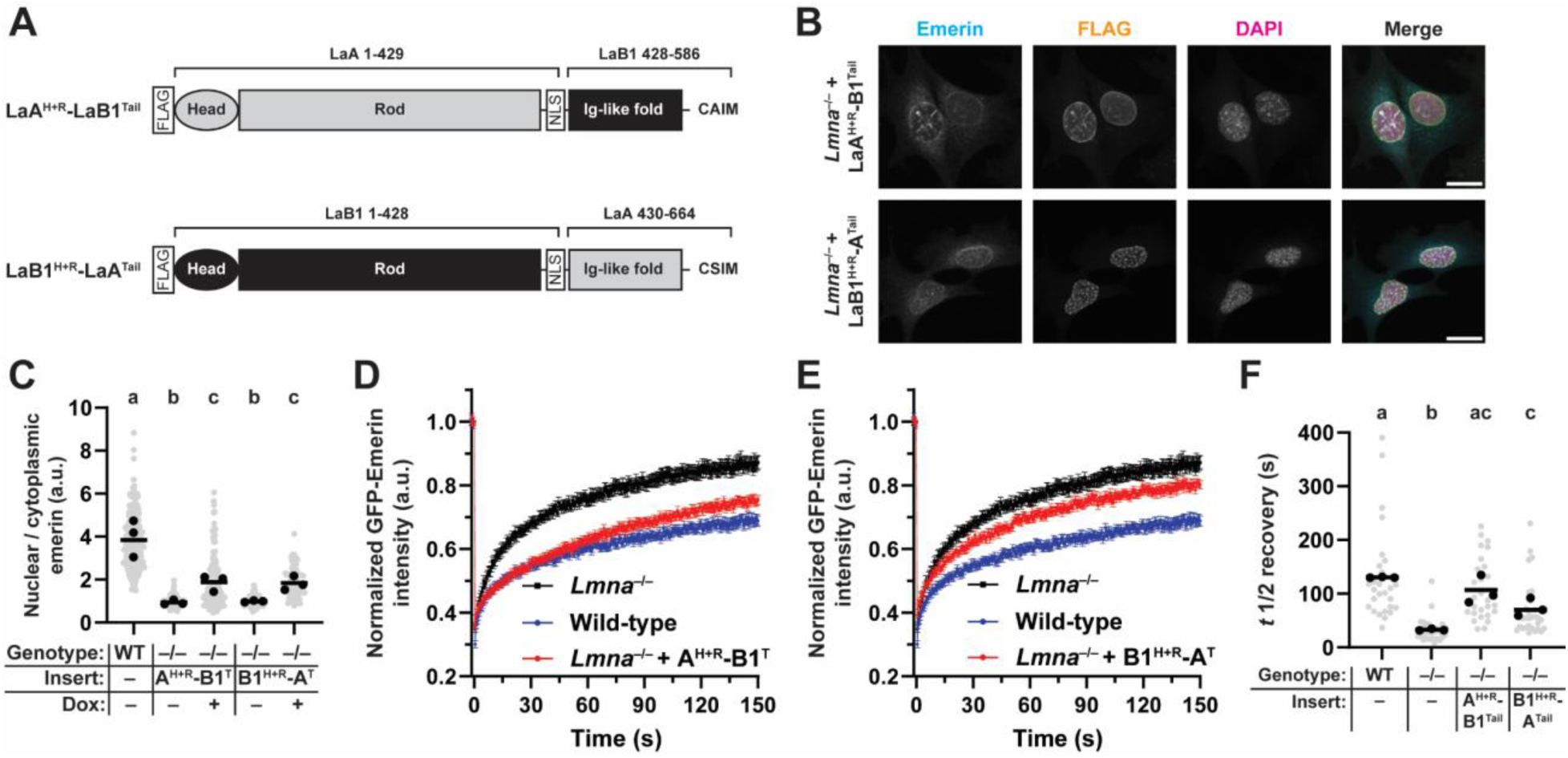
Chimeric lamins reveal multiple domains of lamin A involved in emerin interaction. (**A**) Schematic of chimeric lamin constructs. The residues derived from the respective full-length proteins are indicated above each construct, and both chimeras were tagged with an N-terminal FLAG motif to allow for detection by immunofluorescence. (**B**) Representative immunofluorescence image of *Lmna*^−/−^ MEFs expressing chimeric lamins. FLAG signal indicates the expression of the exogenous chimeric lamin. Scale bar = 20 µm. (**C**) Quantification of mean nuclear/cytoplasmic intensity of endogenous emerin in *Lmna*^−/−^ MEFs expressing chimeric lamins. (**D-E**) FRAP curves of mean GFP-emerin intensity in *Lmna*^−/−^ MEFs expressing LaA^Head+Rod^-B1^Tail^ chimera (**D**) or LaB1^Head+Rod^-A^Tail^ chimera (**E**). Data from wild-type and parental *Lmna*^−/−^ MEFs is shown for reference. (**F**) Quantification of *t*½ median recovery as in Figure 2. Grey points indicate measurements from individual cells, black points indicate replicate means, and bars indicate overall means. Sets of points with the same letter above them are not significantly different, whereas different letters indicate *p* < 0.05, based on one-way ANOVA with Tukey’s multiple comparison test.

#### Permanent anchoring of LaA to the INM enhances retention of emerin at the INM

If the assembly of the LaA network is important for its ability to interact with emerin in cells, we speculated that increased anchoring of LaA to the INM enhances its interaction with emerin compared to wild-type LaA. During LaA posttranslational processing, the C-terminal CaaX motif (C = cysteine, a = aliphatic, X = any amino acid) of prelamin A is a target for farnesylation. After nuclear import and trafficking to the INM, pre-LaA normally undergoes a proteolytic cleavage of the last 15 amino acids, resulting in mature LaA that is not farnesylated (Sinensky *et al*., 1994). The segmental aging disease Hutchinson-Gilford Progeria Syndrome (HGPS) is predominantly caused by a silent mutation in *LMNA* that reveals a cryptic splice site acceptor and results in an in-frame 50 amino acid deletion in the tail of LaA, abolishing the recognition site for the proteolytic cleavage of the C-terminus (De Sandre-Giovannoli *et al*., 2003; Eriksson *et al*., 2003; Glynn and Glover, 2005). The resulting mutant LaA, called progerin, remains permanently farnesylated and aberrantly anchored to the INM, where it forms an extremely stable network, even more persistent than wild-type LaA (Wang *et al*., 2012), leading to nuclear abnormalities such as extensive wrinkling of the nuclear envelope, DNA damage, perturbed nuclear membrane protein localizations and NPC assembly, altered nuclear import/export, mRNA splicing and chromatin organization/regulation, and increased stiffness of the nucleus (Verstraeten *et al*., 2008; Booth *et al*., 2015).

Prior work indicates that progerin binds to emerin in vitro and that progerin may have an improved ability to interact with emerin compared to wild-type LaA (Wu *et al*., 2014). To determine whether enhanced targeting of LaA to the INM influenced its ability to interact with emerin, we generated constructs to allow for expression of FLAG-tagged progerin or truncated progerin (progerin^Tail^) in *Lmna*^−/−^ MEFs, analogous to the LaA^Tail^ construct described previously (Figure 6A). Expression of full-length progerin completely restored the normal nuclear localization of emerin (Figure 6B-C), in agreement with prior reports that progerin is capable of binding to emerin despite lacking 50 amino acids in the tail domain (Wu *et al*., 2014). In contrast with LaA^Tail^, which failed to affect the subcellular localization of emerin, progerin^Tail^ significantly improved the nuclear localization of endogenous emerin to wild-type levels (Figure 6B-C), demonstrating that the targeting of LaA^Tail^ to the nuclear periphery via its farnesyl group was sufficient to promote an enhanced interaction with emerin. Expression of full-length progerin in *Lmna*^−/−^ MEFs significantly reduced the mobility of GFP-emerin following photobleaching, and the *t*_½_ recovery was significantly larger than in wild-type cells (Figure 6D, F). However, in contrast to full-length progerin, expression of the progerin^Tail^ construct in *Lmna*^−/−^ MEFs had no effect on the mobility of GFP-emerin after photobleaching (Figure 6D-F), despite being able to rescue the subcellular distribution of emerin. The progerin^Tail^ construct lacks a lamin rod domain, which is crucial for the formation of lamin dimers and higher-order filaments (Kapinos *et al*., 2010; Zwerger *et al*., 2015). Thus, although the progerin^Tail^ construct can bind to emerin and rescue its nuclear envelope localization, progerin^Tail^ is unable to form a stable network to firmly anchor emerin at the INM and prevent lateral diffusion. Taken together, these results indicate that enhanced anchoring of LaA to the INM can strengthen the interaction with emerin, but that anchoring emerin in place at the INM requires assembly of an A-type lamin filament network.

**Figure 6:**
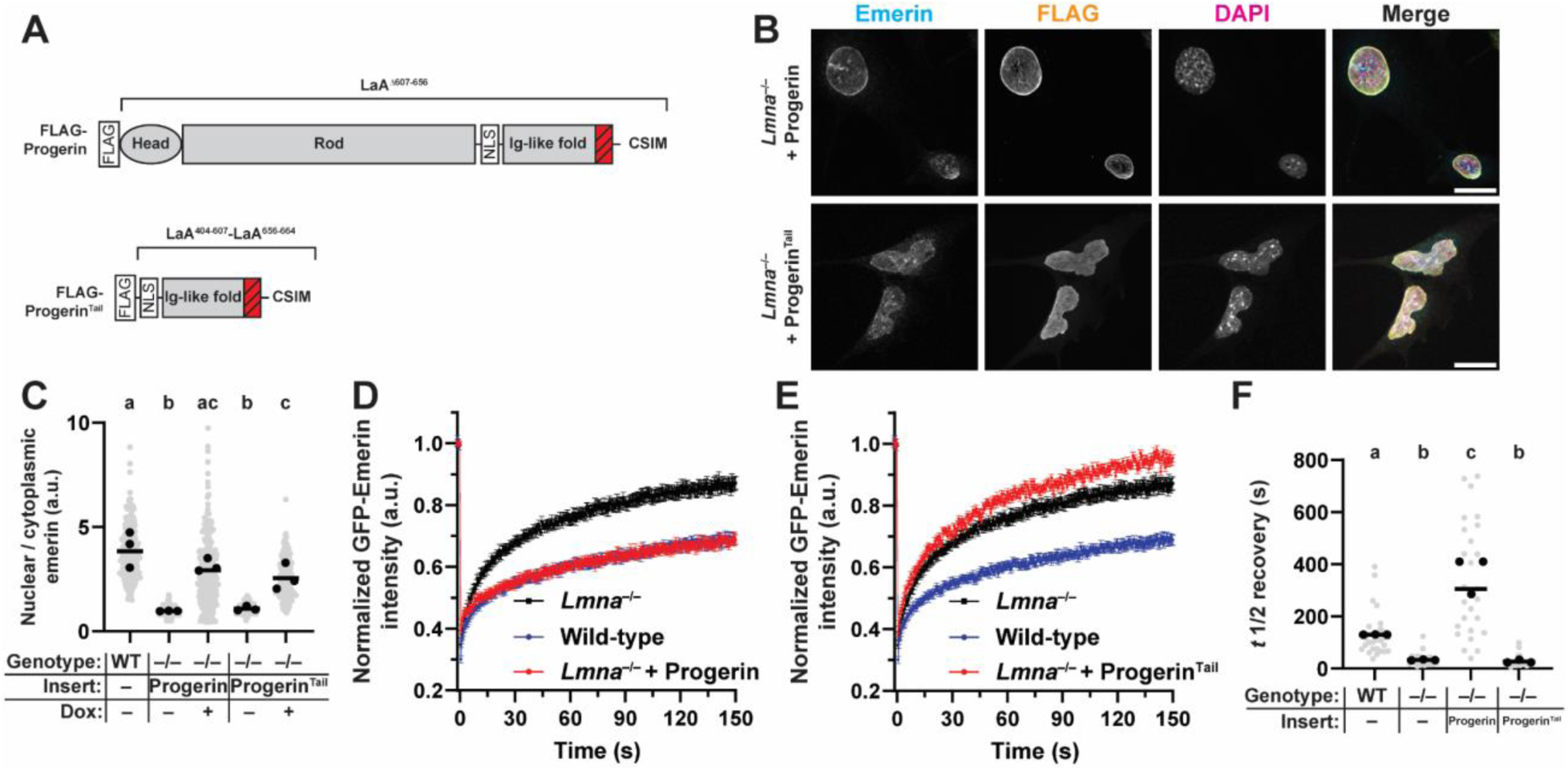
Anchoring LaA to the INM enhances retention of emerin at the nuclear membranes. (**A**) Schematic representation of expression constructs for progerin and progerin^Tail^. Progerin contains a 50 amino acid deletion in the tail of LaA (Δ607-656), indicated in the diagram with the red hash marks. Both constructs contain an N-terminal FLAG tag to allow for detection with anti-FLAG antibodies. (**B**) Representative immunofluorescence images of *Lmna*^−/−^ MEFs expressing progerin or progerin^Tail^. Scale bar = 20 µm. (**C**) Quantification of mean nuclear-to-cytoplasmic fluorescence intensity of emerin in *Lmna*^−/−^ MEFs expressing progerin constructs. (**D-E**) FRAP curves of mean GFP-emerin fluorescence intensity in *Lmna*^−/−^ MEFs expressing progerin (**D**) or progerin^Tail^ (**E**). Data from wild-type and parental *Lmna*^−/−^ MEFs are shown for reference. (**F**) Quantification of *t*½ median recovery of data presented in D-E. Note the increased recovery time of GFP-emerin in cells expressing progerin relative to wild-type MEFs. Grey points indicate measurements from individual cells, black points indicate replicate means, and bars indicate overall means. Sets of points with the same letter above them are not significantly different, whereas different letters indicate *p* < 0.05, based on one-way ANOVA with Tukey’s multiple comparison test.

### Preventing assembly of A-type lamin filaments abolishes anchorage of emerin to the INM

To further explore the requirement of lamin filament assembly on emerin anchorage to the INM, we expressed designed ankryin repeat proteins (DARPins) that specifically bind to LaA/C and interfere with the incorporation of LaA/C into the nuclear lamina in live cells (Zwerger *et al*., 2015). Importantly, the LaA/C-specific DARPin binds to aa 113-140 of LaA, which does not overlap with the LaA emerin-interacting domains in the LaA linker or Ig-fold of the lamin monomer. If emerin requires an intact LaA network to be anchored to the INM, expression of a DARPin that perturbs LaA/C filaments, (DARPin LaA_2), in *Lmna*^−/−^ MEFs expressing LaA should prevent emerin anchorage to the INM. This effect would be detectable by faster recovery time of GFP-emerin following photobleaching. As a control, we expressed a DARPin previously shown to be unable to bind to LaA (DARPin E3.5) and that does not affect LaA filaments (Zwerger *et al*., 2015). *Lmna*^−/−^ MEFs expressing DARPin E3.5 exhibited normal LaA localization following induction of expression (i.e., LaA was enriched at the nuclear periphery). In contrast, cells expressing DARPin LaA_2 exhibited an increased pool of nucleoplasmic LaA signal and a reduction in the peripheral, lamina-associated, pool of LaA (Figure 7A-B). Co-expression of DARPin LaA_2 and LaA in *Lmna*^−/−^ MEFs resulted in a loss of emerin anchorage at the INM and significantly increased mobility of GFP-emerin after photobleaching compared to cells expressing the control DARPin E3.5 (Figure 7C-D). Similar results were obtained with *Lmna*^−/−^ MEFs expressing LaC, where expression of DARPin LaA_2, but not DARPin E3.5, resulted in increased nucleoplasmic LaC localization and abolished the anchoring of emerin to the INM (Figure 7E-H). These results indicate that disrupting the incorporation of either exogenously expressed LaA or LaC into the lamina prevents emerin from being anchored to the INM in *Lmna*^−/−^ MEFs.

**Figure 7:**
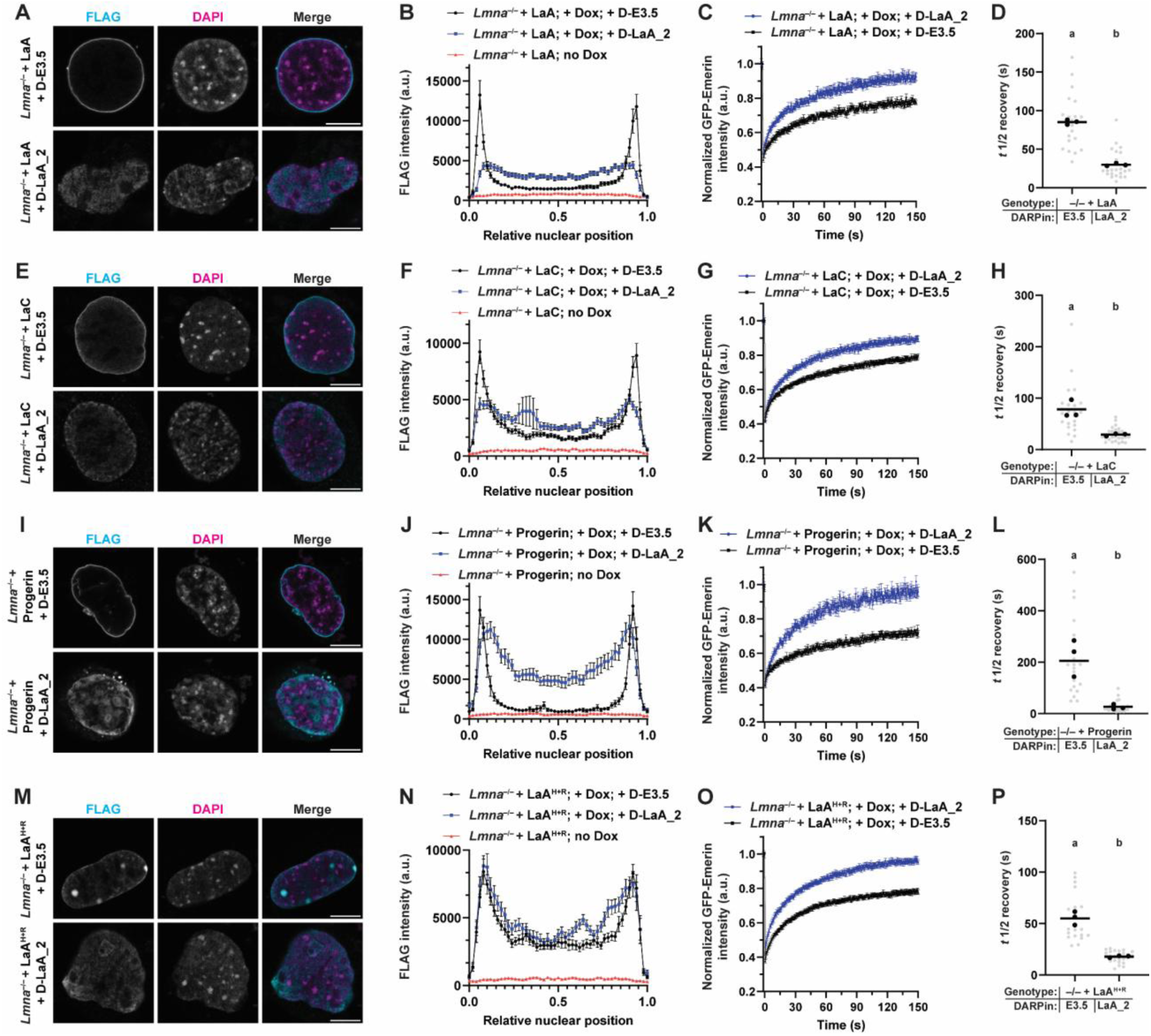
Assembly of A-type lamins into the nuclear lamina is required to anchor emerin at the INM. (**A**) Representative images of *Lmna*^−/−^ MEFs expressing FLAG-LaA and a control DARPin (D-E3.5) or a DARPin that interferes with LaA/C assembly in vivo (D-LaA_2). In both conditions, LaA expression was induced by doxycycline (dox) addition to the media simultaneously with DARPin transfection. Cells were allowed to express the DARPin for 24 h to allow at least one round of cell division to occur in the presence of the DARPin. (**B**) FLAG-LaA fluorescence intensity profile measurements taken by drawing a line through the midplane of the nucleus. Results are displayed as mean ± s.e.m. (**C**) FRAP curves of mean GFP-emerin fluorescence intensity in *Lmna*^−/−^ MEFs expressing LaA and either DARPin E3.5 or DARPin LaA_2. (**D**) Quantification of *t*_½_ median recovery for the data shown in C. Note the increased recovery time of GFP-emerin in cells expressing D-LaA_2. (**E**) Representative images of *Lmna*^−/−^ MEFs expressing LaC and either DARPin E3.5 or DARPin LaA_2. (**F**) FLAG-LaC intensity profile measurements taken by drawing a line through the midplane of the nucleus. Results are shown as mean ± s.e.m. (**G**) FRAP curves of mean GFP-emerin fluorescence intensity in *Lmna*^−/−^ MEFs expressing LaC and either DARPin E3.5 or DARPin LaA_2. (**H**) Quantification of *t*_½_ median recovery of data shown in G. Note the increased recovery time of GFP-emerin in cells expressing D-LaA_2. (**I**) Representative images of *Lmna*^−/−^ MEFs expressing progerin and either DARPin E3.5 or DARPin LaA_2. (**J**) FLAG-progerin fluorescence intensity profile measurements taken by drawing a line through the midplane of the nucleus. Results are shown as mean ± s.e.m. (**K**) FRAP curves of mean GFP-emerin intensity in *Lmna*^−/−^ MEFs expressing progerin and either DARPin E3.5 or DARPin LaA_2. (**L**) Quantification of *t*_½_ median recovery of data presented in K. Note the increased recovery time of GFP-emerin in cells expressing D-LaA_2. (**M**) Representative images of *Lmna*^−/−^ MEFs expressing LaA^H+R^ and either DARPin E3.5 or DARPin LaA_2. (**N**) FLAG-LaA^H+R^ intensity profile measurements taken by drawing a line through the midplane of the nucleus. Results are shown as mean ± s.e.m. (**O**) FRAP curves of mean GFP-emerin fluorescence intensity in *Lmna*^−/−^ MEFs expressing LaA^H+R^ and either DARPin E3.5 or DARPin LaA_2. (**P**) Quantification of *t*_½_ median recovery of data presented in panel O. Note the increased recovery time of GFP-emerin in cells expressing D-LaA_2. Grey points indicate measurements from individual cells, black points indicate replicate means, and horizontal bars indicate overall means. Sets of points with the same letter above them are not significantly different, whereas different letters indicate *p* < 0.05, based on unpaired two-tailed *t*-tests.

We performed similar experiments with *Lmna*^−/−^ MEFs expressing progerin. Expression of the DARPin LaA_2 was sufficient to prevent proper incorporation of progerin into the lamina, resulting in displacement of progerin into the nucleoplasm (Figure 7I-J) and significantly reduced anchorage of emerin to the INM, resulting in faster recovery of GFP-emerin after photobleaching (Figure 7K-L). Finally, expression of DARPin LaA_2 in *Lmna*^−/−^ MEFs expressing LaA^Head+Rod^ significantly increased the mobility of GFP-emerin following photobleaching (Figure 7M-P). Collectively, these results indicate that assembly of A-type lamins into a peripheral filamentous lamin network is a prerequisite for effective anchoring of emerin to the INM, because when lamin filaments were disrupted using DARPins, neither LaA nor LaC was able to anchor emerin at the INM.

As an alternative approach to inhibit LaA filament formation, we expressed a phosphomimetic LaA construct (LaA^PM^: S22D/S392D), which mimics two key sites of LaA phosphorylation that occur at the onset of mitosis to disassemble the lamina and has been shown previously to have aberrant nuclear localization and increased solubility compared to wild-type LaA (Kochin *et al*., 2014). As expected, following expression in *Lmna*^−/−^ MEFs, LaA^PM^ displayed increased nucleoplasmic signal compared to LaA^WT^ (Supplemental Figure 5B). However, LaA^PM^ remained somewhat enriched at the nuclear periphery, which differs from the effect seen following DARPin expression, where peripheral localization of LaA is almost completely abolished (compare Supplemental Figure 5B with Figure 7A). This suggests that the phosphomimetic mutations are unable to completely prevent LaA assembly, whereas the DARPins block lamina assembly more completely. When expressed in *Lmna*^−/−^ MEFs, LaA^PM^ was able to restore the nuclear localization of endogenous emerin to the same degree as wild-type LaA (Supplemental Figure 5B-C), but LaA^PM^ had an impaired ability to anchor GFP-emerin to the nuclear envelope compared to LaA^WT^ based on FRAP experiments (Supplemental Figure 5D-E). These data support our model that lamin assembly is important for LaA to tightly anchor emerin to the nuclear envelope, since interference with LaA assembly via phosphomimetic mutations or DARPins impaired the ability of LaA to restrict emerin’s lateral diffusion.

## Discussion

In this study, we aimed to understand the requirements for anchoring emerin at the INM by A-type lamins, including the role of specific lamin domains. A table summarizing the experiments performed with the various lamin constructs and their ability to rescue emerin localization and anchoring in *Lmna*^−/−^ MEFs is shown in Figure 8. Our studies identified that A-type lamins contain *two* separate emerin-interaction domains: the previously described region involving the Ig-like fold domain (Samson *et al*., 2018), and a newly identified region (aa 394-420) comprising the linker between the LaA rod and Ig-like fold. LaB1 lacks homology to LaA/C within the LaA linker and Ig-like fold regions, and thus the overexpression of LaB1 cannot compensate for the loss of A-type lamins in *Lmna*^−/−^ MEFs. We present a model where two LaA/C domains act additively to interact with emerin and highlight that effective anchoring of emerin to the INM requires not only binding of emerin to A-type lamins, but also the formation of a network of A-type lamin filaments at the nuclear lamina.

**Figure 8:**
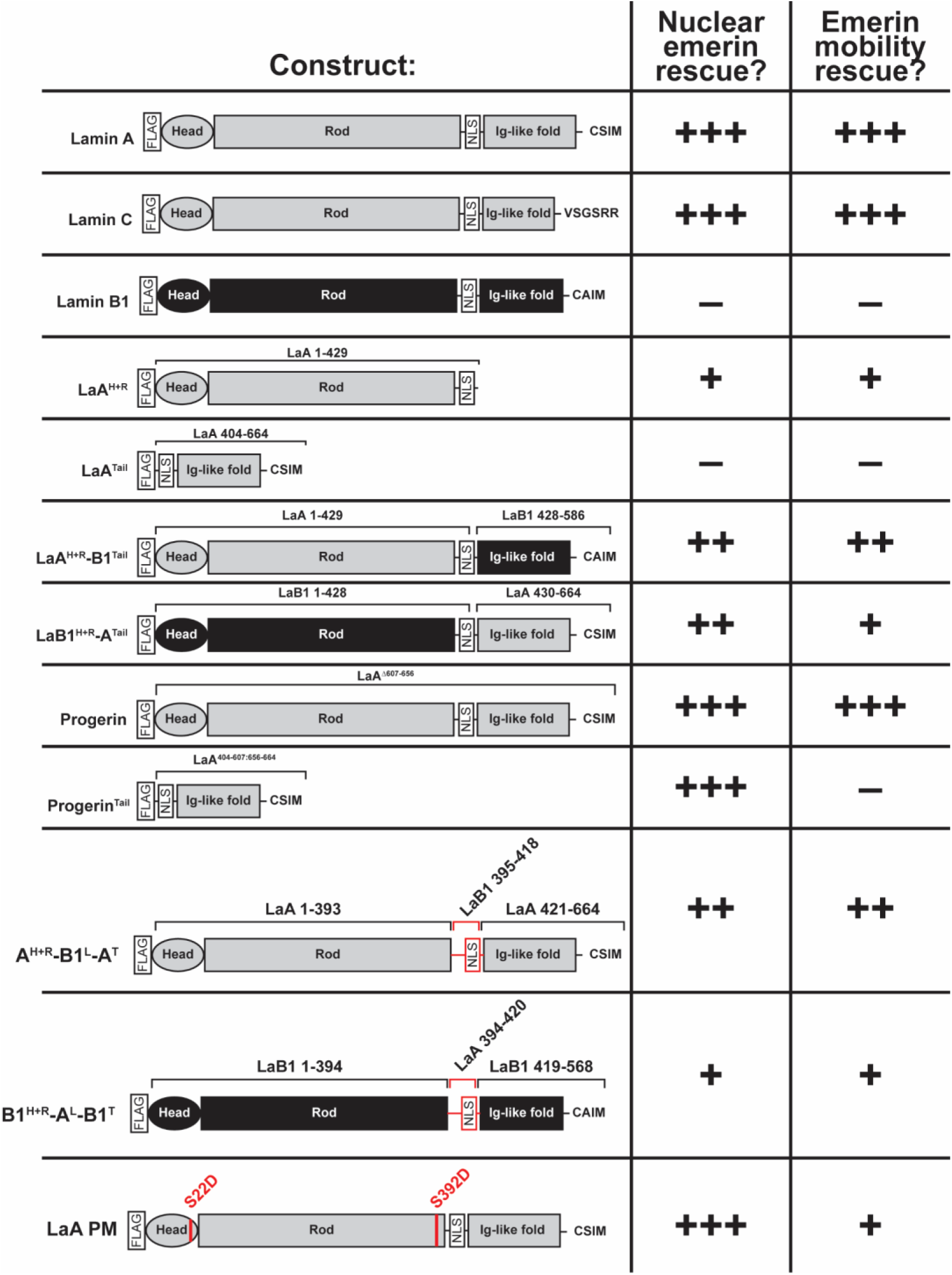
Summary of emerin rescue experiments. Table showing the magnitude of rescued nuclear emerin localization and FRAP mobility of GFP-emerin when the indicated constructs were expressed in *Lmna*^−/−^ MEFs. The number of “+” symbols represents the degree of rescue achieved by each construct; “–” indicates no rescue relative to *Lmna*^−/−^ MEFs not expressing any A-type lamins.

Our experimental data demonstrating the importance of multiple lamin domains in emerin binding and anchorage may help explain why EDMD-causing mutations in *LMNA* tend to occur in either coil 2 (where they disrupt lamin dimerization) or in the Ig-like fold (where they disrupt the interaction of A-type lamins with emerin) (Holt *et al*., 2003; Bertrand *et al*., 2020). In particular, LaA mutations in the LaA rod that do not directly perturb the emerin interaction domains in the linker and Ig-like fold can still result in EDMD, which can be explained by our experimental evidence that effective anchorage of emerin to the nuclear envelope requires not only the presence of an emerin interaction domain, but also an intact lamin rod domain (evidenced by the lack of rescue with the LaA^Tail^, for example) that is essential for lamin dimerization and filament formation, and an assembled LaA network (shown by the DARPin and LaA^PM^ experiments).

Notably, neither the previously established emerin-interacting domain within the LaA/C Ig-like fold nor the newly identified emerin-interacting domain within the LaA/C linker was able to achieve complete rescue relative to full-length wild-type LaA/C. In our experiments using lamin chimeras containing the individual emerin-interacting domains, i.e., the LaA^Head+Rod^-B1^Tail^ and reciprocal constructs, each construct achieved a partial rescue in emerin localization and mobility that amounted to about half of the rescue achieved by full-length LaA. These findings suggest that the two emerin-interacting domains of A-type lamins act additively to achieve proper emerin localization and function, although we cannot exclude that they might also have some synergistic effects on emerin binding. Additive contributions of multiple lamin domains to proper function of lamins within cells may be relevant for other laminopathies caused by mutations in *LMNA*. For example, in *LMNA*-related dilated cardiomyopathy, disease-causing mutations are located across the length of the *LMNA* gene (Worman, 2012). *LMNA* mutations may result in impaired abilities to form stable lamina networks, thus causing fragile nuclei prone to rupture and damage (En *et al*., 2024; Zuela-Sopilniak *et al*., 2024; Pavlov *et al*., 2025), or may lead to disrupted abilities to bind other nuclear proteins like emerin and transcriptional regulators, leading to altered gene expression and manifesting in disease (Ho *et al*., 2013).

Based on the UMD-LMNA database (umd.be/LMNA), several disease-associated human mutations have been described within the LaA linker region (395-420), a subset of which are specific to LaA (R399C/H, G411D, G413C, V415I) (Florwick *et al*., 2017), although none have been specifically linked to EDMD. In general, mutations in the LaA linker region may be poorly tolerated or lethal in the human population and therefore would not be expected to present as muscular dystrophy. In particular, mutations that interfere with the NLS (aa 417-420) or the ability of the NLS to be recognized by importin subunits would likely have deleterious effects in vivo, as the mutant lamin would no longer be imported into the nucleus and accumulate on ER membranes before being released into the cytoplasm (Wu *et al*., 2014).

### Limitations of our current study

All exogenous lamin constructs used in this work possessed an N-terminal FLAG tag, which has been previously observed to slightly impair interaction between lamins and emerin (Odell and Lammerding, 2024). The use of an epitope tag was necessary to assess the localization and relative expression levels of each construct. It was also critical to allow the co-immunoprecipitation experiments to be performed using a single antibody to pull down different bait proteins. By intentionally using a small tag and adding the same tag to all expression constructs, we believe that any biases are introduced uniformly and controlled for by the comparison to the corresponding controls. Another limitation is that the experiments used human lamins expressed in mouse cells, and we thus cannot fully account for potential species-specific differences between the lamin homologs. Nonetheless this limitation does not preclude us from making direct comparisons between the different exogenous lamin constructs.

Future work will be needed to determine whether the interaction between the LaA linker and emerin is direct, or if it requires intermediate binding proteins such as BAF, which has been implicated in mediating the interaction between emerin and the LaA/C Ig-like fold (Samson *et al*., 2018). Although we cannot exclude that BAF might mediate in part some of the interaction between LaA/C and emerin, ample evidence exists for direct interaction between LaA and emerin, including in vitro binding assays using recombinant proteins (Clements *et al*., 2000; Berk *et al*., 2014), which indicates that direct binding between LaA and emerin is possible in the absence of BAF. Additionally, our results with Man1-GFP suggest that the ability of LaA to tightly anchor emerin to the nuclear membrane is not shared with all LEM domain proteins, indicating BAF-independent mechanisms. It also remains to be determined whether the two emerin-interacting domains of LaA/C can bind to the same emerin molecule or if they recognize separate emerin domains. Previous work using surface plasmon resonance has estimated the stoichiometry of emerin bound to each LaA protein between 2:1 and 3:1 (Clements *et al*., 2000), which may reflect emerin’s ability to self-interact (Berk *et al*., 2014) and/or point towards LaA being able to bind multiple emerin molecules simultaneously.

The existence of a novel emerin-interaction domain within the LaA linker is strongly supported by prior in vitro binding experiments between LaA and emerin that included the residues identified in the LaA linker (aa 394-420) in their recombinant LaA fragments. For example, both Sakaki and colleagues and Berk and colleagues demonstrate emerin binding using LaA C-terminal constructs beginning at aa 385 and extending until aa 664 and 646, respectively (Sakaki *et al*., 2001; Berk *et al*., 2014). To our knowledge, our work is the first study to investigate constructs that separate the LaA/C linker residues from the Ig-like fold, which may explain why the existence of multiple emerin-binding sites of LaA/C has not been reported previously.

Our findings have broad relevance for fundamental cell biology, as emerin is expressed in many tissues, and the proper subcellular localization of emerin is crucial for its ability to interact with transcriptional regulators and participate in mechanoresponsive signaling (Lammerding *et al*., 2005; Berk *et al*., 2013). Proper interaction between A-type lamins and emerin is particularly important in muscle, as *LMNA* mutations that disrupt emerin subcellular localization or anchoring are likely to perturb emerin signaling and contribute to the pathology of EDMD. Collectively, the present study improves our understanding of the requirements for LaA/C to anchor emerin at the INM in mammalian cells and explains why the interaction between lamins and emerin is specific to A-type rather than B-type lamins.

## Methods

### Cell culture

Wild-type and *Lmna*^−/−^ MEFs were a kind gift of Colin Stewart (Sullivan *et al*., 1999). Triple lamin knockout MEFs were a kind gift from Stephen Young and Loren Fong and have been described in depth previously (Chen *et al*., 2018, 2021). Cells were maintained in DMEM supplemented with 10% FBS and 1% penicillin/streptomycin in a humidified incubator set to 37°C. Cells were passaged at 80-90% confluency and routinely checked for mycoplasma contamination by PCR. For stable genetic manipulations, the PiggyBac transposase system was used as described previously (Odell *et al*., 2025). Antibiotic selection was performed using Puromycin at 3 µg/ml for at least 1 week, or until cells in a non-transformed well had all died. To induce expression of the exogenous lamin, doxycycline was added at 100 ng/ml for 24 h prior to experiments.

### Genetic construct information

Expression constructs for full-length human lamins, truncated lamins, and a subset of the chimeric lamins have been described previously (Odell *et al*., 2025). The progerin^Tail^, linker-swapped, and phosphomimetic lamin constructs were generated by Gibson cloning. A table of primers used for PCR amplification of these fragments is described in Supplemental Table 1. Inserts were cloned into the pPB-rtTA-hCas9-puro-PB backbone (Wang *et al*., 2017) following digestion with NheI and AgeI. Successful cloning of each construct was determined by Sanger sequencing. For PiggyBac transposition, plasmids containing the desired insert were co-transfected into cells with a plasmid encoding a hyperactive transposase (2:1 vector plasmid: hyperactive transposase plasmid) using Mirus TransIT-X2 transfection reagent according to manufacturer’s instructions.

### Immunofluorescence

Cells were seeded on fibronectin-coated 12 mm glass coverslips overnight, then doxycycline (100 ng/ml) was added for 24 hours to induce expression of the exogenous lamin. Cells were fixed in 4% paraformaldehyde in PBS for 15 min at room temperature, followed by three 5-min washes with IF wash buffer containing 0.2% Triton X-100, 0.25% Tween 20 and 0.3% BSA in PBS. Cells were blocked in 3% BSA in PBS for 1 h, then primary antibodies were added for 1 h in blocking buffer at room temperature or overnight at 4°C. Primary antibodies used were: anti-FLAG (Sigma F7425, 1:1000), and anti-emerin (Leica NCL-emerin, 1:500). DAPI was added 1:500 in PBS for 15 minutes to stain DNA. Secondary antibodies used were Alexa Fluor 488 or 568-conjugated donkey anti mouse or rabbit-IgG antibodies (Invitrogen) diluted 1:250 in 3% BSA in PBS. Coverslips were mounted on glass slides using Mowiol and kept in the dark until imaging. Confocal immunostaining images were acquired on a Zeiss LSM900 series confocal microscope with airyscan module using a 40× water immersion objective. The optimal z-slice size was automatically determined using Zen Blue (Zeiss) software. Airy units for images were set between 1.5 and 2.5.

### Fluorescence recovery after photobleaching (FRAP)

For FRAP experiments, 1 × 10^5^ cells were seeded on a fibronectin coated 35 mm glass-bottom dish the day before transfection. Transient transfection of GFP-emerin (Östlund *et al*., 1999) or Man1-GFP (Wu *et al*., 2002) was achieved using Mirus TransIT-X2 transfection reagent according to manufacturer’s instructions. To transiently express DARPins for FRAP studies, plasmids encoding mCherry-DARPin E3.5 or LaA_2 (Zwerger *et al*., 2015) were co-transfected with the GFP-emerin plasmid; only cells with mCherry signal were used for FRAP experiments. 100 ng/ml doxycycline was added simultaneously with transfection reagent to induce expression of the desired lamin construct. Cells were allowed to express GFP-emerin, Man1-GFP, or DAPRins and the exogenous lamin for 24 hours prior to FRAP experiments. Photobleaching was performed using Zeiss LSM 900 confocal microscope with a 40 × water immersion objective (NA 1.20) using the “iterative bleaching” function. The focus was set to the central z-position for each cell, such that the GFP-emerin nuclear rim was visible. A ROI at the nuclear envelope was drawn manually for each cell. Three frames were acquired before photobleaching, then GFP-emerin was bleached using 100% 488 laser power at maximum scan speed (corresponding to a pixel dwell time of 0.687 µs) for 10 iterations within the ROI, followed by image acquisition every 0.5 s for 150 s following photobleaching. Efforts were made to select cells for photobleaching with comparable GFP-emerin expression levels, and similarly sized ROIs for bleaching were drawn across conditions.

### FRAP analysis

Image analysis was performed using a FIJI macro available upon request. Mean fluorescence intensity of GFP-emerin within the bleached regions was determined for all frames of the timelapse. The mean fluorescence intensity of a background region devoid of cells was subtracted from the fluorescence intensity measurements, and data were normalized to the total cell fluorescence intensity at each time point to control for general photobleaching as a result of continuous imaging. Data were then normalized to the average pre-bleach intensity to yield the normalized intensity over time within the bleached region. On average, the ROI bleaching depth was around 70%, meaning that around 70% of the signal was lost immediately after photobleaching. FRAP data from 20-30 cells were averaged to yield the FRAP recovery curves in the manuscript. Data were fit to a single variable exponential curve using Microsoft Excel, and the median *t*_½_ recovery values were calculated according to the methods described previously (Östlund *et al*., 2006).

### Co-Immunoprecipitation and immunoblotting

Cells (5 × 10^5^) were seeded in 10-cm plates, allowed to adhere overnight, then doxycycline was added (100 ng/ml) for 24 hours to induce expression of FLAG tagged lamin. Cells were lysed on ice in high-salt RIPA buffer containing 12 mM sodium deoxycholate, 50 mM Tris-HCl pH 8.0, 750 mM NaCl, 1% (v/v) NP-40 Alternative, 0.1% (v/v) SDS. Lysates were then vortexed for 5 min, sonicated (Branson 450 Digital Sonifier) for 30 s at 36% amplitude, and centrifuged at 4°C for 10 min at 14,000 g. For each co-IP, 20 µL of anti-FLAG beads (Sigma M8823) were washed twice in high-salt RIPA buffer (300 µL) to equilibrate the beads. 1% of the whole cell lysate was stored at −80°C, and the remainder was mixed with the equilibrated anti-FLAG beads and the total volume was brought up to 1 ml. Immune complexes were allowed to form overnight at 4°C with gentle agitation. Proteins bound to the beads were then washed 5 × with co-IP wash buffer (50 mM Tris-HCl pH 8.0, 300 mM NaCl, 0.3% (v/v) Triton X). Bound proteins were eluted from beads using 100 µL low-pH elution buffer (0.1 M Glycine pH 3.0) for 10 minutes with orbital rotation at 40 rpm. Eluates were immediately quenched with 20 µL 1M Tris (pH 8.0) to neutralize the samples. Protein concentration was determined by Pierce BCA Protein Assay Kit (ThermoFisher A65453) and equal amounts of protein from 1% input and IP eluates were mixed with 4× Laemmli buffer (Biorad 1610747). Samples were denatured by boiling for 3 min, loaded onto 4–12% Bis-Tris gels (Invitrogen NP0322), run for 1.5 h at 100 V, then transferred for 1 h at 16 V onto PVDF membrane.

Membranes were blocked for 1 h in blocking buffer containing 3% BSA in Tris-buffered saline plus 1% Tween 20. Primary antibodies used: anti-FLAG (Sigma F7425, 1:2000), anti-emerin (Leica NCL-emerin, 1:1000). Secondary antibodies used were: Licor IRDye 800CW Donkey anti-Rabbit IgG (926-32213; 1:5000). Secondary antibodies were added for 1 h at room temperature in blocking buffer, followed by three 10-min washes. Membranes were imaged using the Odyssey Licor scanner, and then cropped and brightness and contrast was adjusted using Image Studio Lite (version 5.2) software. Band intensities were determined using the automatic band detection feature of Image Studio Lite. To compare the amount of emerin co-immunoprecipitated with different lamin constructs, we first quantified the intensity of the bands for the co-immunoprecipitated emerin, emerin in the IP input, and immunoprecipitated lamin (based on the FLAG tag). We then divided the levels of the co-immunoprecipitated emerin by the levels of emerin in the input. This fraction was then normalized to the levels of LaA immunoprecipitated in each condition.

### Image analysis

For measurements of nuclear-to-cytoplasmic ratios of endogenous emerin, nuclear emerin immunofluorescence intensity levels were measured using a previously described FIJI macro (Odell *et al*., 2024). Briefly, the image was thresholded based on the DAPI staining to determine the nuclear region of interest (ROI), and the mean fluorescence intensity in the emerin channel within the nucleus was measured. Then, mean cytoplasmic emerin fluorescence intensity levels were obtained by manually drawing a ROI adjacent to the corresponding nucleus for each cell. Mean nuclear emerin fluorescence intensity was divided by mean cytoplasmic emerin fluorescence intensity for each cell. Colocalization analysis was performed using JACoP (Just Another Colocalization Plugin) plugin (Bolte and Cordelières, 2006), following manual thresholding to identify nuclear regions. Intensity profile measurements were performed using a FIJI macro available on request. Briefly, this macro used the ‘Plot Profile’ feature in FIJI software to measure the FLAG intensity across a line drawn across a z-slice through the center of the nucleus. To account for differences in nuclear size, the intensity profiles are converted into relative nuclear distances by measuring the average intensity in each of 50 equally sized bins. For all analysis, cells on the edges of the image, dead cells, and mitotic cells were excluded manually from the analysis.

### Statistical analysis and figure generation

All analyses were performed using GraphPad Prism. Information on statistical tests used and significance values are present in each figure caption. Our statistical analysis was developed in close consultation with the Cornell Statistical Consulting Unit. Figures were assembled using Adobe Illustrator.

## Acknowledgements

We thank Howard Worman (Columbia University) for generously sharing the GFP-emerin and Man1-GFP constructs and Ohad Medalia (University of Zurich) for sharing the DARPin expression constructs. We thank the Biotechnology Resource Center (BRC) Flow Cytometry Facility (RRID: SCR_021740) and sequencing facility (RRID: SCR_021727) at the Cornell Institute of Biotechnology for their resources and technical assistance. This work was supported by awards from the Volkswagen Foundation (A130142 to J.L.), the National Institutes of Health (R01 HL082792, R01 GM137605, R35 GM153257, and R01 AR084664 to J.L.), the National Science Foundation (URoL2022048 to J.L.), and the Leducq Foundation (20CVD01 and 24CVD03 to J.L.). The content of this manuscript is solely the responsibility of the authors and does not necessarily represent the official views of the National Institutes of Health.

## Notes

### Competing Interest Statement

The authors have declared no competing interest.

### Summary of Updates

The revised manuscript includes additional experimental data, including experiments with cells lacking all endogenous lamins, and experiments with another LEM domain protein, Man1-GFP, that further support the conclusions of the manuscript. The manuscript has been further revised to clarify some statements and add information.

## References

Aebi, U, Cohn, J, Buhle, L, and Gerace, L (1986). The nuclear lamina is a meshwork of intermediate-type filaments. Nature 323, 560–564.

Berk, JM, Simon, DN, Jenkins-Houk, CR, Westerbeck, JW, Grønning-Wang, LM, Carlson, CR, and Wilson, KL (2014). The molecular basis of emerin–emerin and emerin–BAF interactions. J Cell Sci 127, 3956–3969.

Berk, JM, Tifft, KE, and Wilson, KL (2013). The nuclear envelope LEM-domain protein emerin. Nucleus 4, 298–314.

Bertrand, AT, Brull, A, Azibani, F, Benarroch, L, Chikhaoui, K, Stewart, CL, Medalia, O, Ben Yaou, R, and Bonne, G (2020). Lamin A/C Assembly Defects in LMNA-Congenital Muscular Dystrophy Is Responsible for the Increased Severity of the Disease Compared with Emery–Dreifuss Muscular Dystrophy. Cells 9, 844.

Bione, S, Maestrini, E, Rivella, S, Mancini, M, Regis, S, Romeo, G, and Toniolo, D (1994). Identification of a novel X-linked gene responsible for Emery-Dreifuss muscular dystrophy. Nat Genet 8, 323–327.

Bolte, S, and Cordelières, FP (2006). A guided tour into subcellular colocalization analysis in light microscopy. J Microsc 224, 213–232.

Booth, EA, Spagnol, ST, Turi A. Alcoser, and Noel Dahl, K (2015). Nuclear stiffening and chromatin softening with progerin expression leads to an attenuated nuclear response to force. Soft Matter 11, 6412–6418.

Buchwalter, A (2023). Intermediate, but not average: The unusual lives of the nuclear lamin proteins. Curr Opin Cell Biol 84, 102220.

Chen, NY, Kim, P, Weston, TA, Edillo, L, Tu, Y, Fong, LG, and Young, SG (2018). Fibroblasts lacking nuclear lamins do not have nuclear blebs or protrusions but nevertheless have frequent nuclear membrane ruptures. Proc Natl Acad Sci U S A 115, 10100–10105.

Chen, NY, Kim, PH, Tu, Y, Yang, Y, Heizer, PJ, Young, SG, and Fong, LG (2021). Increased expression of LAP2β eliminates nuclear membrane ruptures in nuclear lamin-deficient neurons and fibroblasts. Proc Natl Acad Sci U S A 118, e2107770118.

Clements, L, Manilal, S, Love, DR, and Morris, GE (2000). Direct Interaction between Emerin and Lamin A. Biochem Biophys Res Commun 267, 709–714.

De Sandre-Giovannoli, A, Bernard, R, Cau, P, Navarro, C, Amiel, J, Boccaccio, I, Lyonnet, S, Stewart, CL, Munnich, A, Le Merrer, M, et al. (2003). Lamin A Truncation in Hutchinson-Gilford Progeria. Science 300, 2055–2055.

Dou, Z, Xu, C, Donahue, G, Shimi, T, Pan, J-A, Zhu, J, Ivanov, A, Capell, BC, Drake, AM, Shah, PP, et al. (2015). Autophagy mediates degradation of nuclear lamina. Nature 527, 105–109.

En, A, Bogireddi, H, Thomas, B, Stutzman, AV, Ikegami, S, LaForest, B, Almakki, O, Pytel, P, Moskowitz, IP, and Ikegami, K (2024). Pervasive nuclear envelope ruptures precede ECM signaling and disease onset without activating cGAS-STING in Lamin-cardiomyopathy mice. Cell Rep 43, 114284.

Eriksson, M, Brown, WT, Gordon, LB, Glynn, MW, Singer, J, Scott, L, Erdos, MR, Robbins, CM, Moses, TY, Berglund, P, et al. (2003). Recurrent de novo point mutations in lamin A cause Hutchinson-Gilford progeria syndrome. Nature 423, 293–298.

Florwick, A, Dharmaraj, T, Jurgens, J, Valle, D, and Wilson, KL (2017). LMNA Sequences of 60,706 Unrelated Individuals Reveal 132 Novel Missense Variants in A-Type Lamins and Suggest a Link between Variant p.G602S and Type 2 Diabetes. Front Genet 8.

Glynn, MW, and Glover, TW (2005). Incomplete processing of mutant lamin A in Hutchinson-Gilford progeria leads to nuclear abnormalities, which are reversed by farnesyltransferase inhibition. Hum Mol Genet 14, 2959–2969.

Heitlinger, E, Peter, M, Lustig, A, Villiger, W, Nigg, EA, and Aebi, U (1992). The role of the head and tail domain in lamin structure and assembly: Analysis of bacterially expressed chicken Lamin A and truncated B2 lamins. J Struct Biol 108, 74–91.

Ho, CY, Jaalouk, DE, Vartiainen, MK, and Lammerding, J (2013). Lamin A/C and emerin regulate MKL1–SRF activity by modulating actin dynamics. Nature 497, 507–511.

Holt, I, Östlund, C, Stewart, CL, Man, N thi, Worman, HJ, and Morris, GE (2003). Effect of pathogenic mis-sense mutations in lamin A on its interaction with emerin in vivo. J Cell Sci 116, 3027–3035.

Kapinos, LE, Schumacher, J, Mücke, N, Machaidze, G, Burkhard, P, Aebi, U, Strelkov, SV, and Herrmann, H (2010). Characterization of the Head-to-Tail Overlap Complexes Formed by Human Lamin A, B1 and B2 “Half-minilamin” Dimers. J Mol Biol 396, 719–731.

Koch, AJ, and Holaska, JM (2014). Emerin in health and disease. Semin Cell Dev Biol 29, 95–106.

Kochin, V, Shimi, T, Torvaldson, E, Adam, SA, Goldman, A, Pack, C-G, Melo-Cardenas, J, Imanishi, SY, Goldman, RD, and Eriksson, JE (2014). Interphase phosphorylation of lamin A. J Cell Sci 127, 2683–2696.

Lammerding, J, Hsiao, J, Schulze, PC, Kozlov, S, Stewart, CL, and Lee, RT (2005). Abnormal nuclear shape and impaired mechanotransduction in emerin-deficient cells. J Cell Biol 170, 781–791.

Muchir, A, and Worman, HJ (2007). Emery-Dreifuss muscular dystrophy. Curr Neurol Neurosci Rep 7, 78–83.

Odell, J, Gräf, R, and Lammerding, J (2024). Heterologous expression of Dictyostelium discoideum NE81 in mouse embryo fibroblasts reveals conserved mechanoprotective roles of lamins. Mol Biol Cell 35, ar7.

Odell, J, and Lammerding, J (2024). N-terminal tags impair the ability of lamin A to provide structural support to the nucleus. J Cell Sci 137, jcs262207.

Odell, J, Tang, Y, Ambekar, YS, Kidiyoor, GR, Saadi, H, Woodworth, GF, Holt, LJ, Scarcelli, G, Yu, H, and Lammerding, J (2025). The Deformability of the Mammalian Cell Nucleus is Determined by the Identity of the Lamin Rod Domain. 2025.07.22.666133.

Östlund, C, Ellenberg, J, Hallberg, E, Lippincott-Schwartz, J, and Worman, HJ (1999). Intracellular trafficking of emerin, the Emery-Dreifuss muscular dystrophy protein. J Cell Sci 112, 1709–1719.

Östlund, C, Sullivan, T, Stewart, CL, and Worman, HJ (2006). Dependence of Diffusional Mobility of Integral Inner Nuclear Membrane Proteins on A-Type Lamins. Biochemistry 45, 1374–1382.

Pavlov, DA, Heffler, J, Suay-Corredera, C, Dehghany, M, Shen, KM, Zuela-Sopilniak, N, Randell, R, Uchida, K, Jain, R, Shenoy, V, et al. (2025). Microtubule forces drive nuclear damage in LMNA cardiomyopathy. BioRxiv Prepr Serv Biol, 2024.02.10.579774.

Raharjo, WH, Enarson, P, Sullivan, T, Stewart, CL, and Burke, B (2001). Nuclear envelope defects associated with LMNA mutations cause dilated cardiomyopathy and Emery-Dreifuss muscular dystrophy. J Cell Sci 114, 4447–4457.

Reichart, B, Klafke, R, Dreger, C, Krüger, E, Motsch, I, Ewald, A, Schäfer, J, Reichmann, H, Müller, CR, and Dabauvalle, M-C (2004). Expression and localization of nuclear proteins in autosomal-dominant Emery-Dreifuss muscular dystrophy with LMNA R377H mutation. BMC Cell Biol 5, 12.

Reits, EAJ, and Neefjes, JJ (2001). From fixed to FRAP: measuring protein mobility and activity in living cells. Nat Cell Biol 3, E145–E147.

Sakaki, M, Koike, H, Takahashi, N, Sasagawa, N, Tomioka, S, Arahata, K, and Ishiura, S (2001). Interaction between Emerin and Nuclear Lamins. J Biochem (Tokyo) 129, 321–327.

Samson, C, Petitalot, A, Celli, F, Herrada, I, Ropars, V, Le Du, M-H, Nhiri, N, Jacquet, E, Arteni, A-A, Buendia, B, et al. (2018). Structural analysis of the ternary complex between lamin A/C, BAF and emerin identifies an interface disrupted in autosomal recessive progeroid diseases. Nucleic Acids Res 46, 10460–10473.

Sinensky, M, Fantle, K, Trujillo, M, McLain, T, Kupfer, A, and Dalton, M (1994). The processing pathway of prelamin A. J Cell Sci 107 (Pt 1), 61–67.

Sullivan, T, Escalante-Alcalde, D, Bhatt, H, Anver, M, Bhat, N, Nagashima, K, Stewart, CL, and Burke, B (1999). Loss of a-Type Lamin Expression Compromises Nuclear Envelope Integrity Leading to Muscular Dystrophy. J Cell Biol 147, 913–920.

Vaughan, OA, Alvarez-Reyes, M, Bridger, JM, Broers, JLV, Ramaekers, FCS, Wehnert, M, Morris, GE, Whitfield, WGF, and Hutchison, CJ (2001). Both emerin and lamin C depend on lamin A for localization at the nuclear envelope. J Cell Sci 114, 2577–2590.

Verstraeten, VLRM, Ji, JY, Cummings, KS, Lee, RT, and Lammerding, J (2008). Increased mechanosensitivity and nuclear stiffness in Hutchinson–Gilford progeria cells: effects of farnesyltransferase inhibitors. Aging Cell 7, 383–393.

Wang, G, Yang, L, Grishin, D, Rios, X, Ye, LY, Hu, Y, Li, K, Zhang, D, Church, GM, and Pu, WT (2017). Efficient, footprint-free human iPSC genome editing by consolidation of Cas9/CRISPR and piggyBac technologies. Nat Protoc 12, 88–103.

Wang, Y, Ӧstlund, Cecilia, Choi, Jason C., Swayne, Theresa C., Gundersen, Gregg G., and and Worman, HJ (2012). Blocking farnesylation of the prelamin A variant in Hutchinson-Gilford progeria syndrome alters the distribution of A-type lamins. Nucleus 3, 452–462.

Worman, HJ (2012). Nuclear lamins and laminopathies. J Pathol 226, 316–325.

Wu, D, Flannery, AR, Cai, H, Ko, E, and Cao, K (2014). Nuclear localization signal deletion mutants of lamin A and progerin reveal insights into lamin A processing and emerin targeting. Nucleus 5, 66–74.

Wu, W, Lin, F, and Worman, HJ (2002). Intracellular trafficking of MAN1, an integral protein of the nuclear envelope inner membrane. J Cell Sci 115, 1361–1371.

Zuela-Sopilniak, N, Morival, J, and Lammerding, J (2024). Multi-level transcriptomic analysis of LMNA-related dilated cardiomyopathy identifies disease-driving processes. 2024.06.11.598511.

Zwerger, M, Jaalouk, DE, Lombardi, ML, Isermann, P, Mauermann, M, Dialynas, G, Herrmann, H, Wallrath, LL, and Lammerding, J (2013). Myopathic lamin mutations impair nuclear stability in cells and tissue and disrupt nucleo-cytoskeletal coupling. Hum Mol Genet 22, 2335–2349.

Zwerger, M, Roschitzki-Voser, H, Zbinden, R, Denais, C, Herrmann, H, Lammerding, J, Grütter, MG, and Medalia, O (2015). Altering lamina assembly reveals lamina-dependent and - independent functions for A-type lamins. J Cell Sci 128, 3607–3620.

