## Supplementary Materials for "A-type lamins anchor emerin at the inner nuclear membrane via two independent binding sites"

### Supplemental Materials

#### Supplemental Figures:

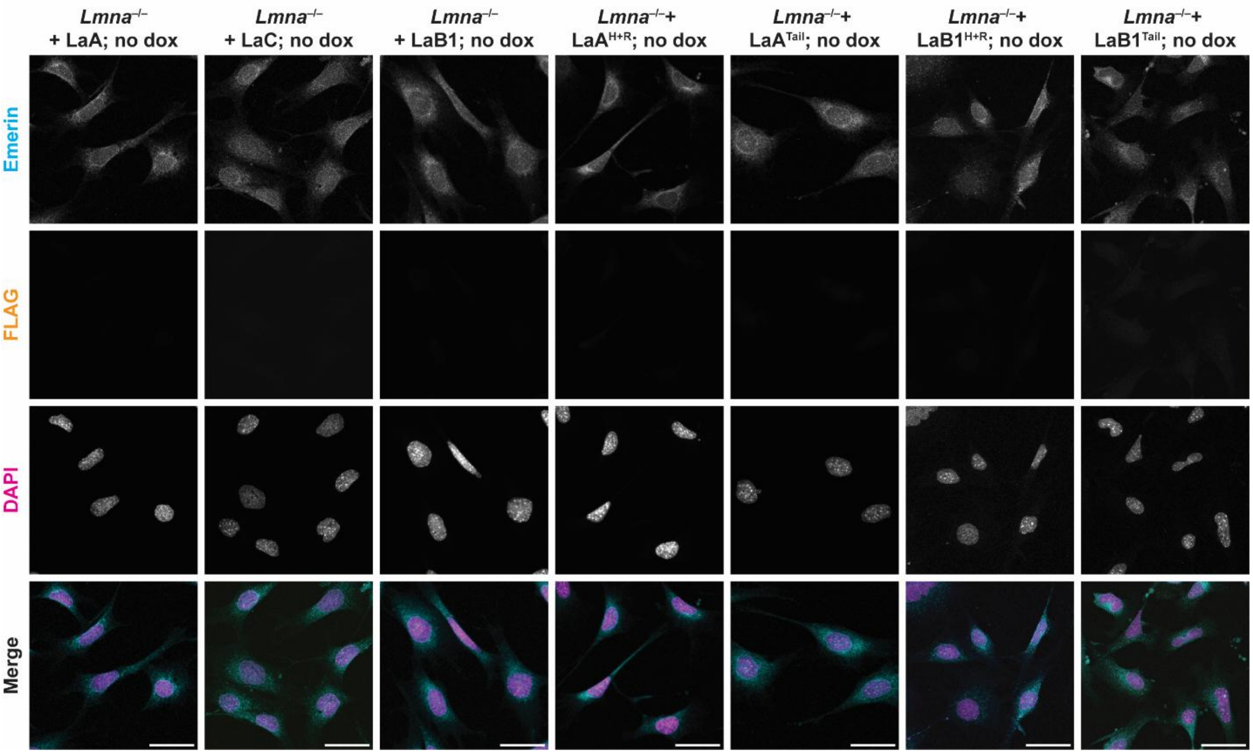


**Supplemental Figure 1: Emerin localization in *Lmna*$^{-/-}$ MEFs without dox induction of exogenous lamins.** Representative images of cells modified with constructs outlined in Figure 1, not treated with dox, and stained for emerin and FLAG. All cells exhibited similar degrees of emerin mislocalization, and there was no leaky expression of the inducible lamin, exhibited by the lack of signal in the FLAG channel. Scale bar = 50 µm.

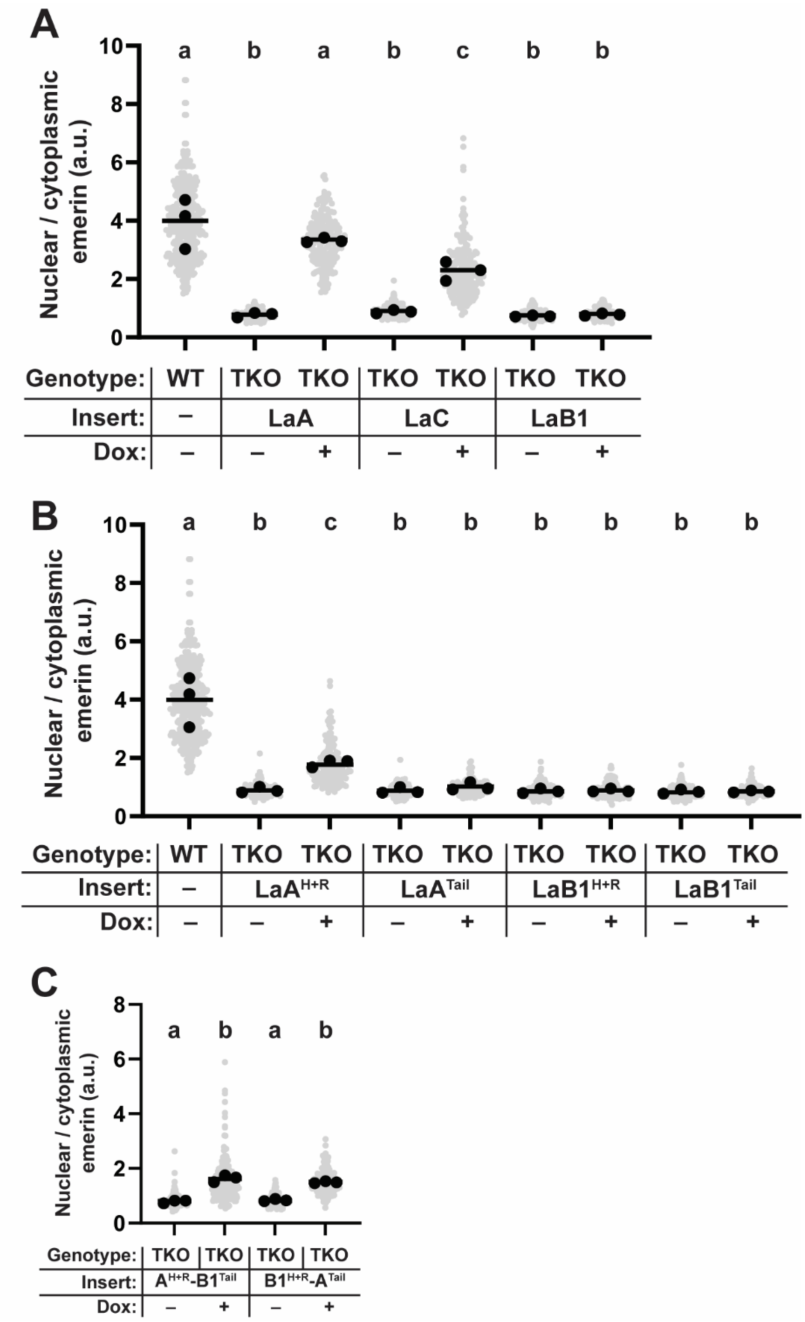


**Supplemental Figure 2: Emerin localization in TKO MEFs expressing different exogenous lamin constructs.** (**A-C**) Quantification of mean nuclear/cytoplasmic fluorescence intensity of immunofluorescently labeled endogenous emerin in TKO MEFs expressing full-length lamins (**A**), truncated lamins (**B**), or chimeric lamins (**C**). Grey points indicate measurements from individual cells, black points indicate replicate means, and bars indicate overall means. Sets of points with the same letter above them are not significantly different, whereas different letters indicate $p < 0.05$, based on one-way ANOVA with Tukey's multiple comparison test.

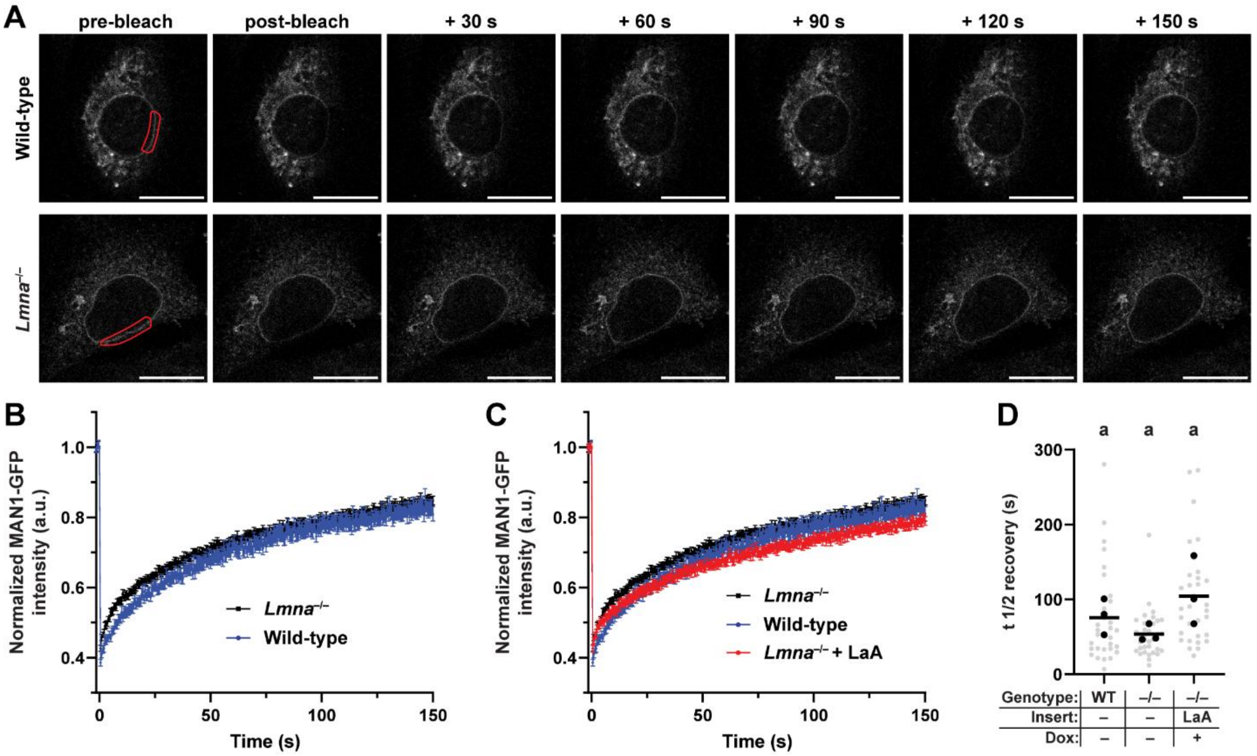


**Supplemental Figure 3: Man1-GFP FRAP in wild-type and *Lmna*$^{-/-}$ MEFs.** (**A**) Representative frames of a time series showing FRAP experiment on Man1-GFP transiently expressed in wild-type (WT) or *Lmna*$^{-/-}$ MEFs. The target bleach area is outlined in red, and the time in seconds following bleach is indicated above each set of images. Scale bar = 10 µm. (**B**) Recovery curves of Man1-GFP following photobleaching in wild-type and *Lmna*$^{-/-}$ MEFs. (C) Recovery curve of Man1-GFP in *Lmna*$^{-/-}$ MEFs expressing LaA. Data from wild-type and *Lmna*$^{-/-}$ cells are shown for reference. (**D**) Quantification of $t_{½}$ median recovery for the data presented in panels B-C. Grey points indicate measurements from individual cells, black points indicate replicate means, and bars indicate overall means. Sets of points with the same letter above them are not significantly different, whereas different letters indicate $p < 0.05$, based on one-way ANOVA with Tukey's multiple comparison test.

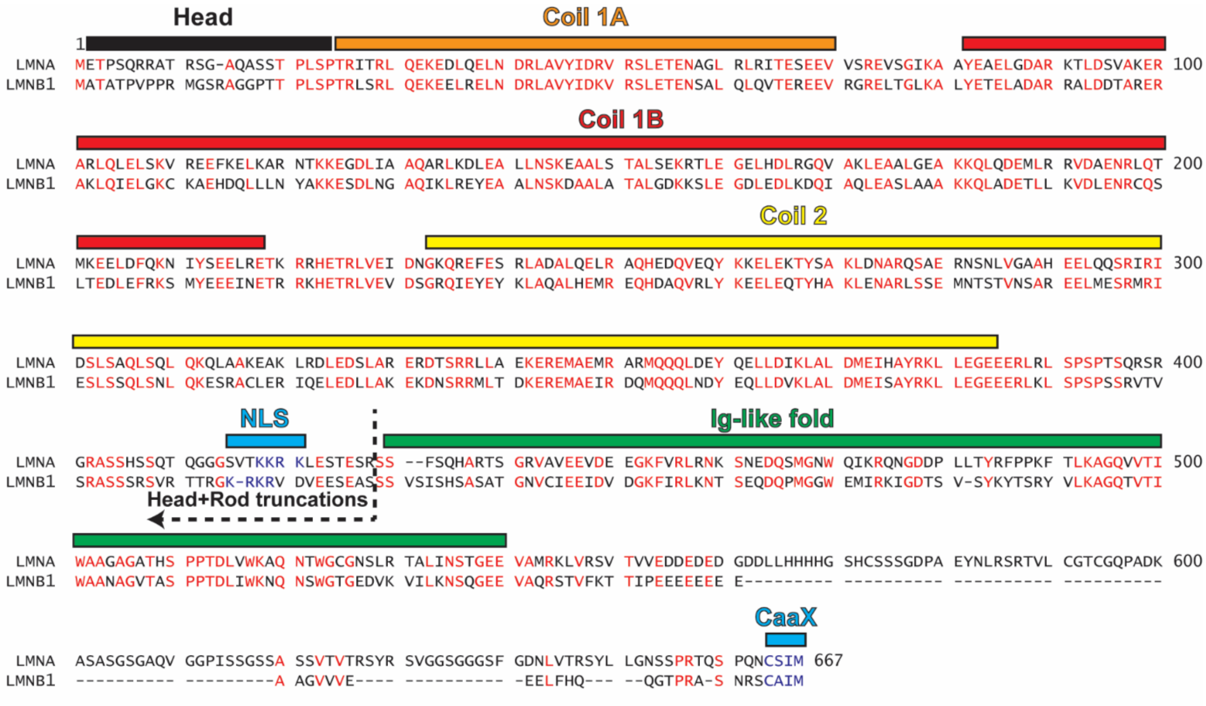


**Supplemental Figure 4: Sequence alignment between human Lamin A and Lamin B1.** Identical residues between the two proteins are colored red. The major structural domains of the lamin proteins are indicated. NLS: nuclear localization signal; Ig-like fold: Immunoglobulin-like fold. The cutoff of residues found in the rod truncations is also indicated by the dashed line next to the NLS.

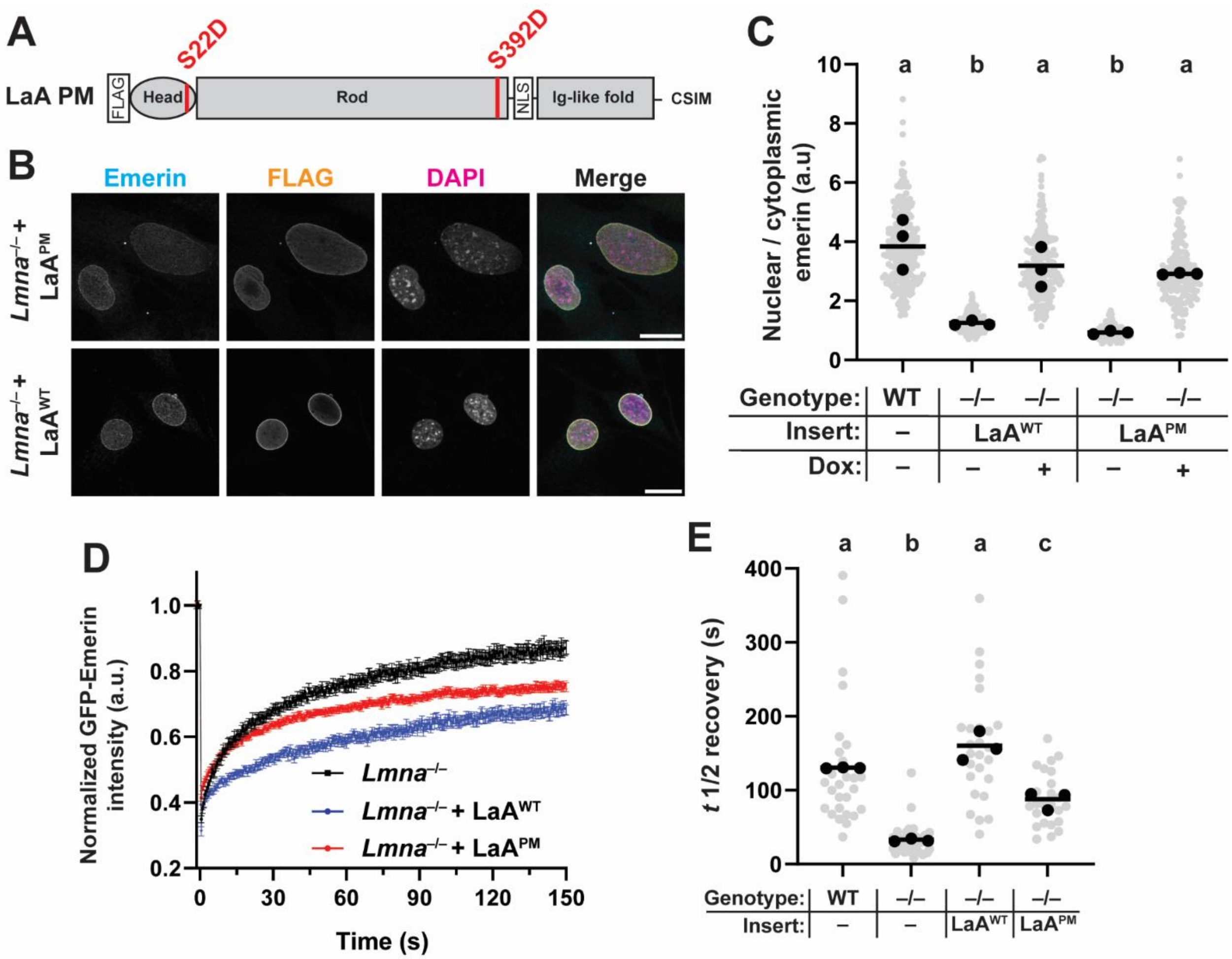


**Supplemental Figure 5: Emerin localization and mobility in *Lmna*$^{-/-}$ MEFs expressing phosphomimetic LaA.** (**A**) Schematic depiction of the phosphomimetic LaA mutations (S22D/S392D) used in the dox inducible expression constructs. (**B**) Representative immunofluorescence images of *Lmna*$^{-/-}$ MEFs expressing either LaA$^{PM}$ or LaA$^{WT}$, each with the same N-terminal FLAG tag. Note the increased nucleoplasmic signal in the FLAG channel for the LaA$^{PM}$ construct. Scale bar = 20 µm. (**C**) Quantification of mean nuclear/cytoplasmic fluorescence intensity of immunofluorescently labeled endogenous emerin in *Lmna*$^{-/-}$ MEFs expressing the indicated lamin construct. (**D**) FRAP curves of mean GFP-emerin intensity in *Lmna*$^{-/-}$ MEFs expressing LaA$^{PM}$. Data from wild-type and *Lmna*$^{-/-}$ MEFs expressing LaA$^{WT}$ is shown for reference. (**E**) Quantification of $t_{½}$ median recovery as in Figure 2. Grey points indicate measurements from individual cells, black points indicate replicate means, and bars indicate overall means. Sets of points with the same letter above them are not significantly different, whereas different letters indicate $p < 0.05$, based on one-way ANOVA with Tukey's multiple comparison test.

#### Supplemental Table

| Primer Name | Sequence 5'-3 | Use |
|---|---|---|
| **LaA_IgFold_fwd** | accctcgtaaaggtctagagaccatggactacaaagacgatgacgacaagtctcactcat cccagacac | Cloning ProgerinTail |
| **LaA_rev** | ccgtttaaactcattactaattacatgatgctgcagttc | Cloning ProgerinTail |
| **LaA_rod_fwd** | accctcgtaaaggtctagagaccatggactacaaagacgatgacg | Cloning LaA w/ B1 Linker |
| **LaB1_linker_rev** | GTACACTACGACTTGAGGATGCTCGGGATACTGTCAC ACGGGAAGAAGGGCTGGGGGACAGGCGTA | Cloning LaA w/ B1 Linker |
| **LaB1_linker_fwd** | ATCCTCAAGTCGTAGTGTACGTACAACTAGAGGAAAG CGGAAGAGGCTGGAGTCCACTGAGAGCCg | Cloning LaA w/ B1 Linker |
| **LaA_rev** | ccgtttaaactcattactaattacatgatgctgcagttc | Cloning LaA w/ B1 Linker |
| **LaB1_Rod_fwd** | accctcgtaaaggtctagagaccatggactacaaagacgatgacg | Cloning LaB1 w/ A Linker |
| **LaA_linker_rev** | TGTGTCTGGGATGAGTGAGAGGAAGCACGGCCACGGC TGCGCTGCGAGGTAGGGCTTGGAGACAGCTTCA | Cloning LaB1 w/ A Linker |
| **LaA_linker_fwd** | CTCTCACTCATCCCAGACACAGGGTGGGGGCAGCGTC ACCAAAAAGCGCAAAGTTGATGTGGAAGAATCAGA | Cloning LaB1 w/ A Linker |
| **LaB1_Tail_rev** | CCGTTTAAACTCATTACTAACTACATAATTGCACAGCT TC | Cloning LaB1 w/ A Linker |
| **S22D_Rev** | GCCGGGTGATGCGGGTGGGatcCAGCGGAGTG | Cloning LaA PM |
| **S22D_fwd** | CACTCCGCTGgatCCCACCCGCATCACCCGGC | Cloning LaA PM |
| **S392D_rev** | CGCTGCGAGGTAGGatcGGGGGACAGG | Cloning LaA PM |
| **S392D_fwd** | CCTGTCCCCCgatCCTACCTCGCAGCG | Cloning LaA PM |

**Supplementary Table 1:** List of primers used for Gibson cloning the indicated lamin constructs.
